# Dendrites with specialized glial attachments develop by retrograde extension using SAX-7 and GRDN-1

**DOI:** 10.1101/801761

**Authors:** Elizabeth R. Cebul, Ian G. McLachlan, Maxwell G. Heiman

## Abstract

Dendrites develop elaborate morphologies in concert with surrounding glia, but the molecules that coordinate dendrite and glial morphogenesis are mostly unknown. *C. elegans* offers a powerful model for identifying such factors. Previous work in this system examined dendrites and glia that develop within epithelia, similar to mammalian sense organs. Here, we focus on the neurons BAG and URX, which are not part of an epithelium but instead form membranous attachments to a single glial cell at the nose, reminiscent of dendrite-glia contacts in the mammalian brain. We show that these dendrites develop by retrograde extension, in which the nascent dendrite endings anchor to the presumptive nose and then extend by stretch during embryo elongation. Using forward genetic screens, we find that dendrite development requires the adhesion protein SAX-7/L1CAM and the cytoplasmic protein GRDN-1/CCDC88C to anchor dendrite endings at the nose. SAX-7 acts in neurons and glia, while GRDN-1 acts in glia to non-autonomously promote dendrite extension. Thus, this work shows how glial factors can help to shape dendrites, and identifies a novel molecular mechanism for dendrite growth by retrograde extension.

## INTRODUCTION

Neurons weave their dendritic arbors within a fabric of axons and glia. How dendrites and axons wire together has been intensely studied, but less attention has been given to developmental interactions between dendrites and glia. Nonetheless, the precisely coordinated morphologies of dendrites and glia are striking and play a key role in controlling neural activity. For example, in the cerebellum, Bergmann glia appear to scaffold the growth of Purkinje dendrites (Lordkipanidze and Dunaevsky, 2005) and guide their innervation by specific axon types (Ango et al., 2008). At the fine scale, Bergmann glia wrap individual dendritic spines to control synapse number (Lippman Bell et al., 2010). Similarly, in the hippocampus, dendrites receive Ephrin signals from apposed glial processes that modulate dendritic spine morphology and can also alter synaptic strength (Murai et al., 2003; Nishida and Okabe, 2007; Carmona et al., 2009; Bernardinelli et al., 2014; Perez-Alvarez et al., 2014). These examples have provided a glimpse into the diversity of dendrite-glia contacts in the brain. However, broader exploration of dendrite-glia contacts and the molecular mechanisms that control them has been limited by the difficulty of studying single defined contacts *in vivo* and performing forward genetic screens in mammalian systems. To overcome these challenges, we have turned to the simple nervous system of *C. elegans* as a model for studying specialized dendrite-glia contacts.

*C. elegans* offers several advantages. Each cell can be uniquely identified and has a remarkably stereotyped shape and set of cell-cell contacts; neuron- and glia-specific promoters allow the targeted expression of transgenes in single cells to visualize and manipulate specific neuron-glia contacts in intact living animals; and its powerful genetics permit unbiased screens to identify factors that control neuron-glia interactions. Additionally, *C. elegans* glia share a number of important features with mammalian glia (Mizeracka and Heiman, 2015). This system thus offers a powerful genetic model for dissecting dendrite-glia contacts. Here, we turned our attention to a previously overlooked class of dendrite-glia contacts whose elaborate, tightly apposed membranous structures are reminiscent of the highly coordinated membranous contacts found between astrocytes and dendritic spines at mammalian synapses.

Using super-resolution microscopy, we find that the dendrites of two neuron types, URX and BAG, form specialized attachments with a single glial partner. Using forward genetics, we identify two conserved molecules, SAX-7 and GRDN-1, that are required for the development of these dendrites. SAX-7 is a homolog of the mammalian neuron-glia adhesion molecule L1CAM, and GRDN-1 is a homolog of proteins involved in cell polarity and cytoskeleton organization. Each of these homologs is expressed in the mammalian brain and causes developmental brain disorders when disrupted (Jouet et al., 1994; Vits et al., 1994; Ekici et al., 2010). Using embryonic time-course imaging, we find that URX and BAG dendrites develop by retrograde extension, a type of outgrowth in which the dendrites anchor to their targets and then extend by stretch. SAX-7 and GRDN-1 help to anchor dendrite endings at the nose during embryo elongation. In mutants, dendrites detach from the nose during development and fail to fully extend. SAX-7 acts both in neurons and glia, while GRDN-1 acts in glia to non-autonomously promote URX and BAG dendrite extension. This suggests that dendrite anchoring might occur through early attachments to glia.

Retrograde extension was previously described in the *C. elegans* amphid, which develops as a narrow epithelial tube made up of neurons and glia (Heiman and Shaham, 2009; Fan et al., 2019; Low et al., 2019). In that example, dendrite anchoring requires apical extracellular matrix that lines the lumen of the developing amphid (Heiman and Shaham, 2009; Low et al., 2019). Interestingly, axons of the zebrafish olfactory system also form by retrograde extension, attaching near the surface of the brain while their cell bodies move away (Breau et al., 2017), suggesting that this mode of growth may be shared across species. The results described here define a novel molecular mechanism of retrograde extension used by non-epithelial neurons, and suggest a surprising means by which dendrite morphogenesis can be shaped by interactions with glia.

## RESULTS

### URX and BAG dendrite endings form specialized contacts with a single glial partner

URX and BAG are bilaterally symmetric pairs of sensory neurons in the head that mediate behavioral responses to oxygen and carbon dioxide (Cheung et al., 2004; Gray et al., 2004; Hallem and Sternberg, 2008; Zimmer et al., 2009; Bretscher et al., 2011; Laurent et al., 2015). Each of these neurons extends a single unbranched dendrite to the nose, a distance of ∼100-150 µm in adults (Fig. 1A,C) (Ward et al., 1975). The dendrites reside in distinct nerve bundles (URX, dorsal bundle; BAG, lateral bundle), where they fasciculate closely with other sensory dendrites and several classes of glial cells (Ward et al., 1975). This study focuses on URX and BAG dendrite interactions with one class of glia, the inner labial socket (ILso) glia.

**Fig. 1.**
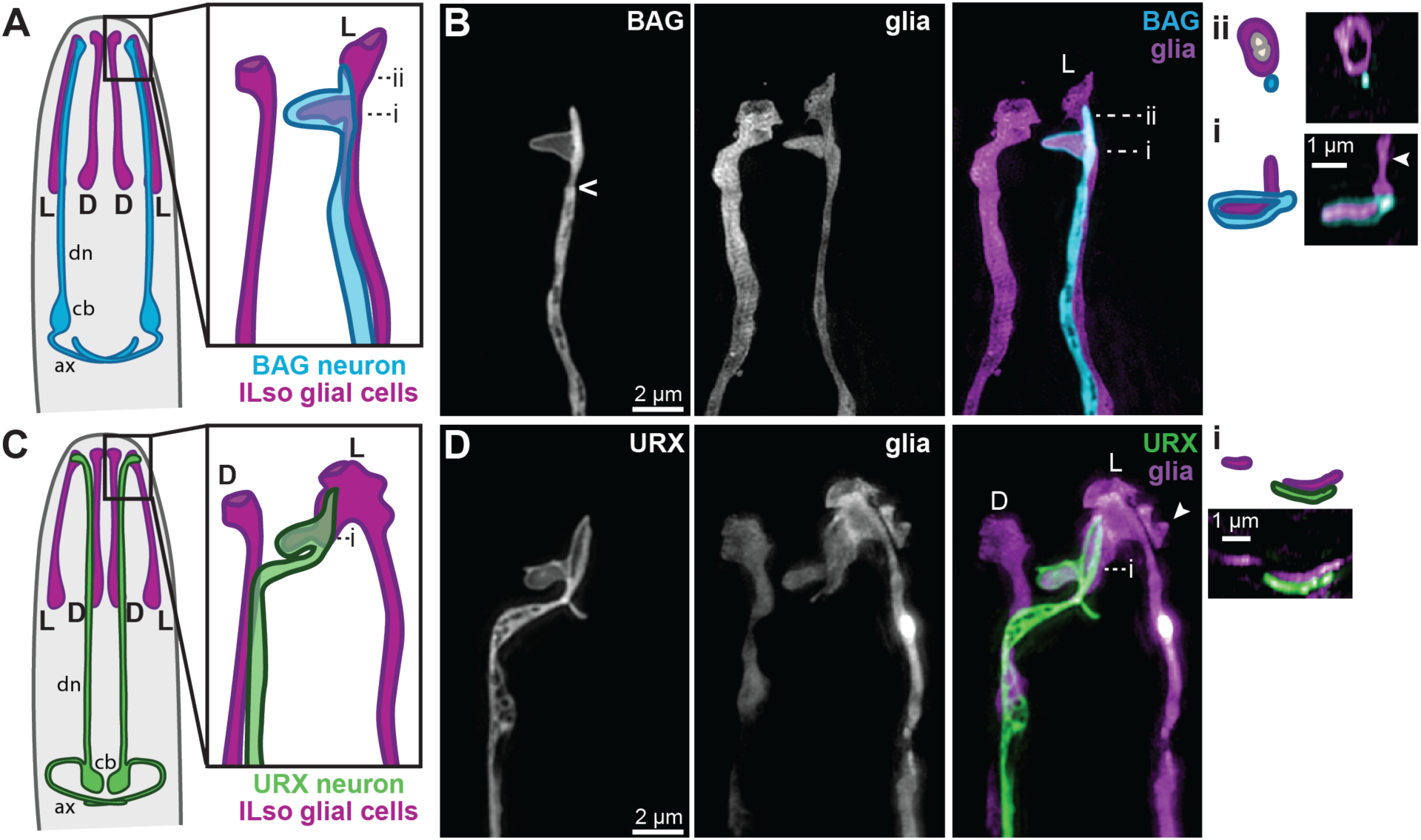
BAG and URX dendrites form specialized contacts with a single glial cell. (A, C) Schematics of a *C. elegans* head (nose is at top) showing BAG (A, blue) or URX (C, green) relative to the inner labial socket glia (ILso, purple). Dorsal (D) and lateral (L) pairs of glia are indicated; an additional ventral pair is not shown. cb, neuronal cell bodies; dn, dendrites; ax, axons. (B,D) Single-wavelength and merged superresolution images of BAG (blue, *flp-17*pro) or URX (green, *flp-8*pro) and ILso glia (purple, *grl-18*pro) acquired with structured illumination microscopy. Caret (<), base of cilium. Insets (Bi) and (Bii) show cross sectional views and schematics of the indicated image planes. (Bi) BAG wraps around a thumb-like protrusion of the glial cell. A second protrusion is visible in this view, arrowhead. (Bii) The ILso glial cell forms a pore through which two unlabeled neurons access the environment; BAG does not enter this pore. (D) The URX dendrite “jumps” from the dorsal to lateral ILso and forms a sheet-like contact with a protrusion from this glial cell; cross section in inset (Di). A second protrusion from the glial cell, likely the BAG-associated thumb, is partly visible, arrowhead.

At the nose, the URX and BAG dendrites terminate in specialized sensory structures containing guanylate cyclases that detect molecular oxygen and carbon dioxide (Smith et al., 2013; Gross et al., 2014). Classical electron microscopy (EM) reconstructions showed that the BAG dendrite terminates in a sensory cilium with a membranous bag-shaped elaboration (hence “BAG”) that wraps around a thumb-like protrusion from the lateral ILso glial cell (Fig. 1A) (Ward et al., 1975; Doroquez et al., 2014). We became interested in this structure because the elaborate wrapping morphology reminded us of glia-dendrite contacts in the mammalian brain. For example, in the cerebellum, Bergmann glia extend elaborate membranous processes that precisely wrap around dendritic spines, structures that are analogous to the BAG sensory cilium (Spacek, 1985; Xu-Friedman et al., 2001; Lippman et al., 2008). Although here the dendrite ending wraps the glial cell, we reasoned that similar mechanisms might underlie coordinated morphogenesis of the dendritic and glial membranes in both cases.

To investigate this structure in intact living animals, we developed tools and approaches for superresolution imaging of BAG and ILso glia. We expressed combinations of soluble, membrane-associated, and ciliary fluorescent markers under the control of cell-specific promoters (see Methods) and used 3D structured illumination microscopy to visualize the cells. We could discern a subtle constriction at the BAG dendrite ending, consistent with the base of the cilium (Fig. 1B, caret), followed by a bag-shaped membranous elaboration ∼1-2 µm in diameter that perfectly wraps a thumb-like protrusion from the lateral ILso glial cell (Fig. 1B; additional examples in Fig. S1A-D). Cross-sectional views confirmed that the BAG cilium wraps around the ILso membrane in this region (Fig. 1Bi). We observed variation in the morphology of this structure across individuals, but the glial cell and the dendritic elaboration always formed precisely interlocking shapes (Fig. S1A-D). In cross-sectional views, we could discern the pore at the ILso glial ending through which dendrites of mechanosensory (IL1) and chemosensory (IL2) neurons protrude to the outside of the animal (Ward et al., 1975); in contrast, the BAG dendrite does not enter this pore and is not directly exposed to the external environment (Fig. 1Bii).

Next, we sought to compare this structure with that of URX. The URX dendrite is not thought to be ciliated, and previous imaging suggested it terminates in a “large bulb” where signaling molecules are localized (Ward et al., 1975; Gross et al., 2014; McLachlan et al., 2018). However, superresolution imaging of the URX dendrite ending revealed a more complex structure (Fig. 1D). As expected, we found that the URX dendrite fasciculates along its length with the dorsal nerve bundle. But, surprisingly, it exits this bundle at the nose and “jumps” over to the lateral nerve bundle (visible in ∼90% of animals, n>40) (Fig. 1D; additional examples in Fig. S1E-H). This contact has not been described previously and suggests that URX can discriminate between the dorsal and lateral ILso glial endings. The URX dendrite does not enter the pore made by the lateral ILso glial cell, but instead forms a membranous sheet-like elaboration that covers another protrusion from this cell (but does not fully wrap it, Fig. 1Di). Thus, the lateral ILso glial cell extends two protrusions, one wrapped by the BAG dendrite ending and the other covered by the URX dendrite ending. The “two-armed” morphology of the lateral ILso glial cell is especially apparent in some views (see Fig. S1A, B, D, H). Remarkably, a similar arrangement is observed in *Pristionchus pacificus*, a species that diverged from *C. elegans* about 100 million years ago (Hong et al., 2019). An intriguing possibility is that this glial cell provides some kind of specialized support for carbon dioxide and oxygen sensation.

Here, we focus on the broad question of how the URX and BAG dendrites develop. Due to their intimate association with the ILso glial cell, we reasoned that their mechanism of development might reveal an interesting role for glia.

### URX and BAG dendrites do not require genes used for amphid dendrite development

We first asked whether URX and BAG extend their dendrites using the molecular mechanism previously described for neurons in the amphid, the largest *C. elegans* sense organ (Heiman and Shaham, 2009). Similar to URX and BAG, each of the 12 amphid neurons has an unbranched dendrite that terminates at the nose, where they display sensory cilia and associate with specific glia (the amphid sheath and socket, Fig. 2A) (Ward et al., 1975). However, unlike URX and BAG, the amphid dendrites form epithelial junctions with glia (Low et al., 2019) and navigate through pores in the glia to directly access the external environment (Fig. 2B). This arrangement resembles the architecture of some mammalian sense organs, like the olfactory epithelium.

**Fig. 2.**
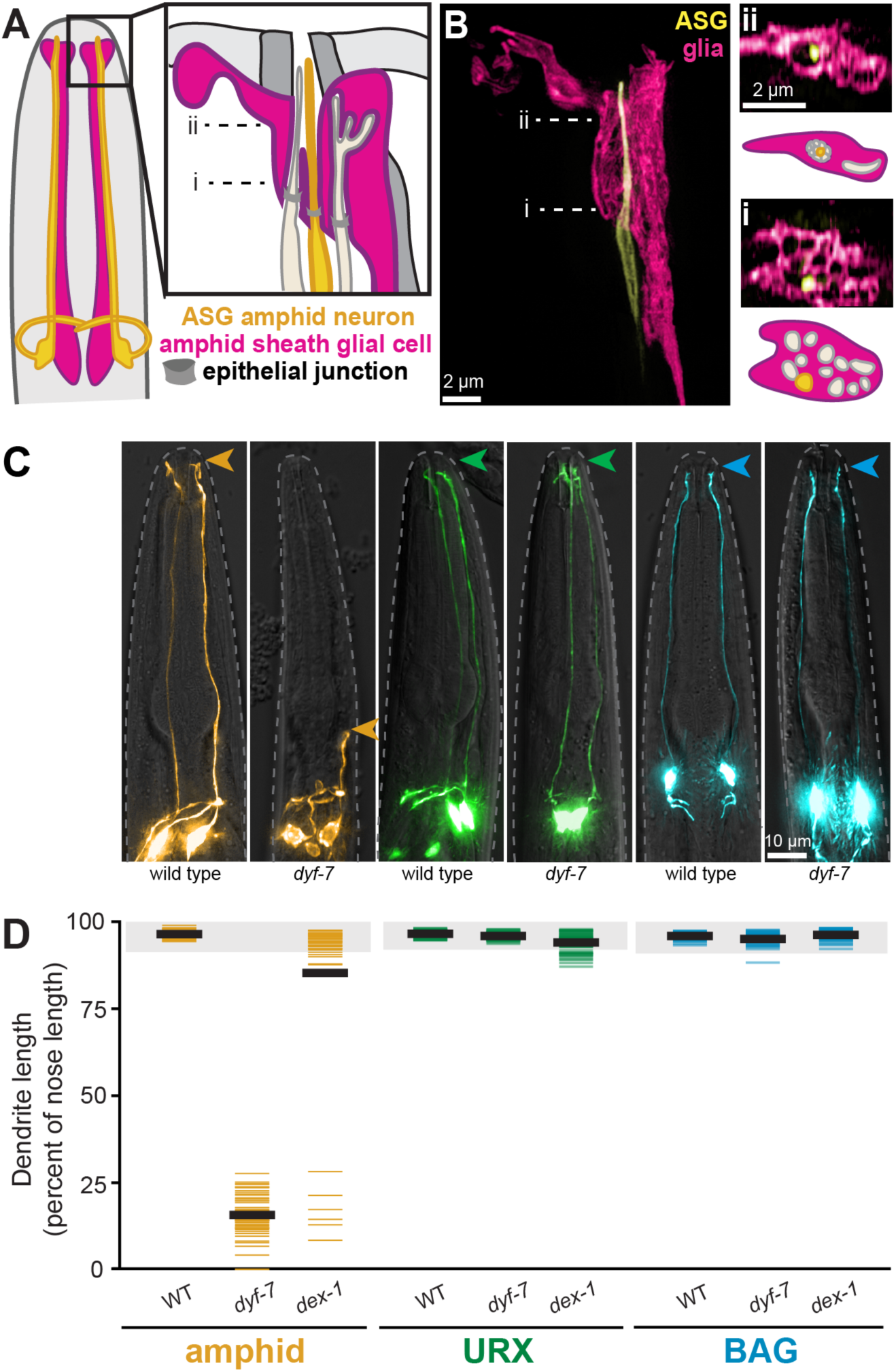
URX and BAG dendrites develop by a mechanism distinct from the amphid. (A) Schematic of a *C. elegans* head showing representative amphid neurons (tan; ASG neuron, yellow) and the amphid sheath glial cell (pink). Most amphid dendrites enter individual channels in the glial cell that converge to a central pore; some terminate in pockets within the sheath. (B) Single-wavelength and merged superresolution images of ASG (yellow, *ops-1*pro) and the amphid sheath (pink, *F16F9.3*pro). Insets i and ii show cross-sectional views and schematics of the indicated image planes. (i) Twelve individual channels in the sheath, one for each amphid dendrite, were previously seen only by EM. ASG can be seen inside one of these channels. (ii) The channels converge distally to a central pore with ASG inside it. (C) Wild-type and *dyf-7* animals expressing markers for an amphid neuron (AWC, yellow, *odr-1*pro), URX (green, *flp-8*pro), or BAG (blue, *flp-17*pro). Arrowheads, dendrite endings. (D) Dendrite lengths in wild-type, *dyf-7*, and *dex-1* animals, expressed as a percentage of the distance from cell body to nose. Colored bars represent individual dendrites (n≥50 per genotype); black bars represent population averages; shaded region represents wild-type mean ± 5 s.d. for each neuron type, which we define as “full length”. All amphid dendrites are affected by *dyf-7* and *dex-1* (Heiman and Shaham, 2009).

Amphid dendrites develop using DYF-7 and DEX-1, which resemble apical extracellular matrix (ECM) proteins that are found in many epithelia but are not widely expressed in the brain (Heiman and Shaham, 2009). During development, DYF-7 and DEX-1 anchor the tips of the developing amphid dendrites at the embryonic nose as the cell bodies migrate away, stretching the dendrites out behind them in a phenomenon called retrograde extension (Heiman and Shaham, 2009). In *dyf-7* or *dex-1* mutants, the dendrite endings detach and the dendrites fail to extend (Heiman and Shaham, 2009). Several other classes of head sensory neurons (CEP, IL, OL) also have dendrite endings that protrude through glial pores to the environment, and all of them also require DYF-7 (Low et al., 2019).

As noted above, URX and BAG dendrites do not protrude through a glial pore and are not directly exposed to the environment. Consistent with these anatomical differences, we found that URX and BAG dendrite development is grossly unaffected in *dyf-7* and *dex-1* mutants (Fig. 2C, D). Some *dex-1* URX dendrites exhibit subtle length defects, possibly due to secondary effects on the IL sense organs. URX and BAG dendrites thus likely develop using different molecular mechanisms than previously studied neurons. This prompted us to try to identify factors required for URX and BAG dendrite development, and to learn if they more closely resemble factors used in the mammalian brain.

### URX and BAG dendrite development requires SAX-7 and GRDN-1

To this end, we performed visual forward genetic screens for mutants in which URX or BAG dendrites fail to fully extend (see Methods). In separate screens focused on each neuron, we isolated eight URX and two BAG dendrite extension mutants (Fig. 3A, URX; 3B, BAG). Each of the mutants exhibits grossly normal body morphology, locomotion, and expression of URX- and BAG-specific markers (*flp-8*, URX; *flp-17*, BAG; *egl-13,* URX and BAG), and ILso glia still extend their processes to the nose (Fig. S2). Together, these observations suggest that the dendrite extension defects are not secondary to pleiotropic abnormalities in morphogenesis, neurodevelopment, or cell fate specification. Interestingly, we found that most of the URX mutants also strongly affect BAG, and vice versa, suggesting that these neurons may employ closely related mechanisms of dendrite extension (Fig. 3). We found that our ten mutants represented two complementation groups, and we proceeded to identify the relevant genes.

**Fig. 3.**
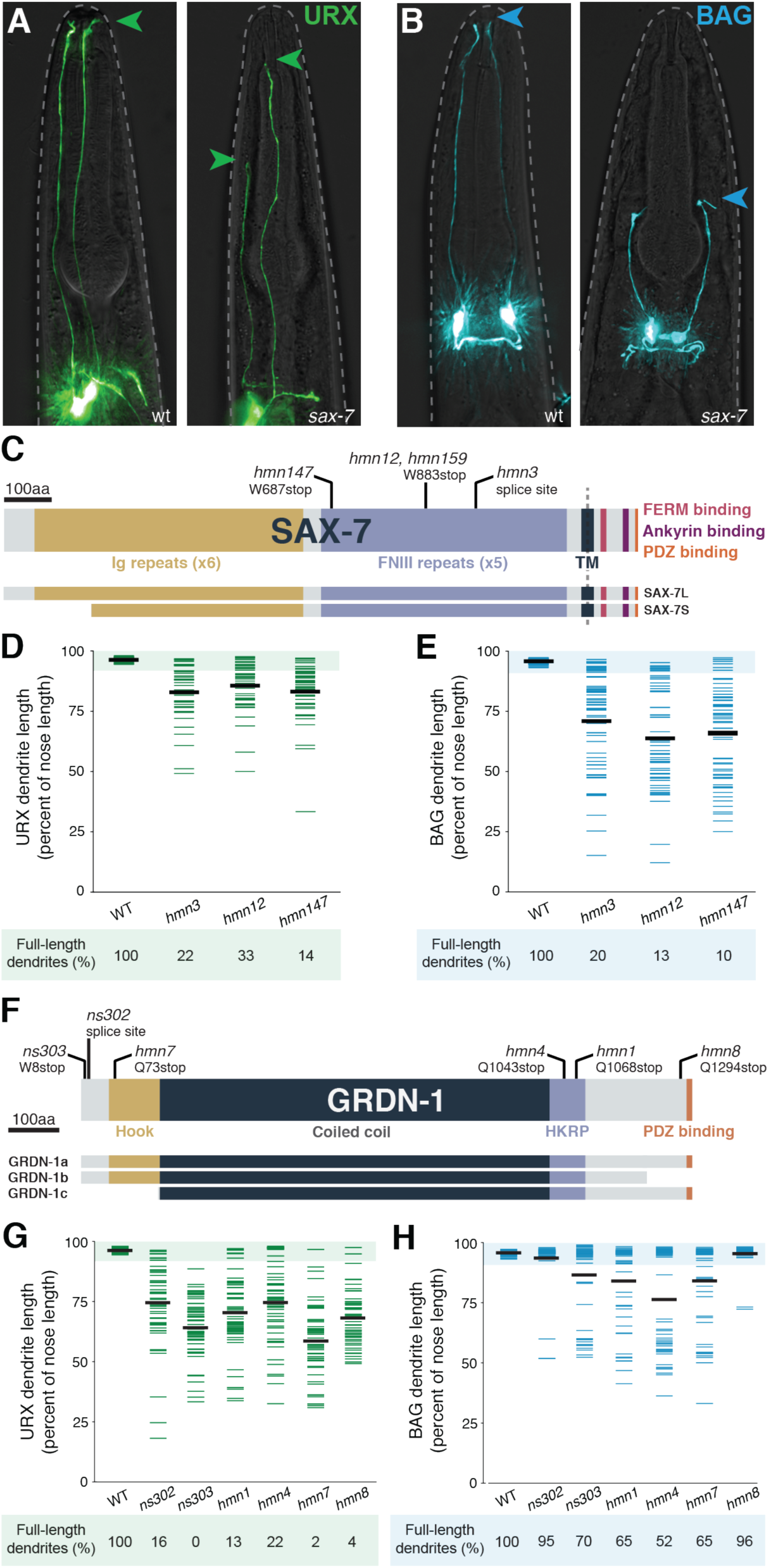
URX and BAG dendrite morphogenesis requires SAX-7 and GRDN-1. (A, B) URX (A, *flp-8*pro) and BAG (B, *flp-17*pro) neurons in wild-type (wt, left) and *sax-7* (right) animals. Arrowheads, dendrite endings. (C, F) Schematics of (C) SAX-7 and (F) GRDN-1 indicating alleles and conserved domains. Ig, immunoglobulin; FNIII, fibronectin type III; TM, transmembrane; HKRP, Hook-related protein. Differences between isoforms are shown below. (D, E, G, H) Quantification of (D, G) URX and (E, H) BAG dendrite lengths, expressed as a percentage of the distance from cell body to nose. Colored bars represent individual dendrites (n≥48 per genotype); black bars represent population averages. Shaded region represents wild-type mean ± 5 s.d. for each neuron type and the percentage of dendrites in this range (“full length dendrites”) is indicated below the plots.

#### SAX-7

Using standard mapping approaches, we found that alleles in the first complementation group (*hmn3, hmn12, hmn147, hmn159*) disrupt *sax-7*, which encodes a transmembrane cell adhesion molecule related to mammalian L1CAM (Fig. 3C) (Zallen et al., 1999; Chen et al., 2001; Sasakura et al., 2005). L1CAM is expressed throughout the brain and plays several roles during development, including in neuron-glia adhesion (Grumet and Edelman, 1984; Stoeckli et al., 1997). Defects in L1CAM lead to X-linked hydrocephalus (L1 Syndrome), a highly variable disorder that can also include spastic paraplegia and intellectual disability (Jouet et al., 1994; Vits et al., 1994; Adle-Biassette et al., 2013). How loss of L1CAM leads to such a broad spectrum of phenotypes remains mysterious. Interestingly, L1CAM is also expressed in some epithelial tissues, but less is known about its roles in this context (Debiec et al., 1998; Nolte et al., 1999; Pechriggl et al., 2017).

In *C. elegans*, SAX-7 affects the development and maintenance of many aspects of nervous system organization, including axon guidance, synapse function, dendrite fasciculation, axon and neuronal cell body positioning, and the spatial patterning of branched dendrite arbors (Zallen et al., 1999; Kim and Li, 2004; Sasakura et al., 2005; Bénard et al., 2012; Dong et al., 2013; Salzberg et al., 2013; Opperman et al., 2015; Yip and Heiman, 2018; Chen et al., 2019; Ramirez-Suarez et al., 2019). In these contexts, SAX-7 has been shown to physically interact with UNC-44/ankyrin, STN-2/γ-syntrophin, the transmembrane protein DMA-1, and the contactin RIG-6 (Zhou et al., 2008; Liu and Shen, 2011; Dong et al., 2013; Salzberg et al., 2013; Kim and Emmons, 2017). However, we found that mutations in these genes do not recapitulate the dendrite extension defects seen in *sax-7* (Fig. S3A).

All of our *sax-7* alleles cause dendrite extension defects in both URX and BAG (Fig. 3D, E), with BAG exhibiting more severe defects in both penetrance and expressivity. For example, in *sax-7(hmn12)*, dendrite extension defects occur in 87% of BAG neurons vs 67% of URX neurons, and the affected BAG dendrites are 59±19% of nose length vs 81±10% for URX. Each of our alleles is predicted to disrupt both the short (SAX-7S) and long (SAX-7L) isoforms, whereas an allele that disrupts only the SAX-7L isoform (*nj53)* has no effect on URX or BAG dendrite length (Fig. S3A) (Sasakura et al., 2005). Surprisingly, two of the alleles we isolated (*hmn12* and *hmn159*) result in an identical nucleotide change despite arising from separate screens focused on URX and BAG, respectively; a previously described allele (*ky146*) also results in the same codon change (Zallen et al., 1999). These alleles are predicted to truncate the protein prior to its transmembrane segment and are thus expected to be functional nulls (Sasakura et al., 2005). We have used *sax-7(hmn12)* throughout this study.

#### GRDN-1

Alleles in the second complementation group (*ns302, ns303, hmn1, hmn4, hmn7,* and *hmn8*) disrupt *grdn-1* (Fig. 3F), which is predicted to encode a soluble cytoplasmic protein related to mammalian Girdin/CCDC88A (hence “*grdn-1*”), CCDC88B, CCDC88C, and other members of the Hook-related protein family (HKRP). Their roles in brain development have not been fully explored, but Girdin can promote long-range neuronal migration by affecting cell adhesion and, separately, can affect synaptic plasticity (Enomoto et al., 2009; Wang et al., 2011; Nakai et al., 2014; Itoh et al., 2016). Loss of Girdin/CCDC88A causes PEHO syndrome, characterized by progressive encephalopathy and optic atrophy (Nahorski et al., 2016). Intriguingly, loss of CCDC88C leads to congenital hydrocephalus similar to that seen in loss of L1CAM/SAX-7 (Ekici et al., 2010; Drielsma et al., 2012; Ruggeri et al., 2018), although no relationship between CCDC88C and L1CAM has been identified previously.

In *C. elegans*, GRDN-1 acts through CHE-10/Rootletin in ciliated sensory neurons to position the ciliary basal body and promote proper cilia development (Nechipurenko et al., 2016). However, we found that neither URX nor BAG dendrite lengths are affected by *che-10* (Fig. S3B), suggesting that the URX and BAG dendrite defects are unlikely to be secondary to the CHE-10-dependent role of GRDN-1 in cilia.

Quantification of our allelic series revealed that most *grdn-1* alleles strongly affect both URX and BAG (Fig. 3G, H); however, in contrast to what we observed with *sax-7*, the *grdn-1* defects are always more severe in URX. Indeed, two alleles (*ns302, hmn8*) produce barely discernible effects on BAG despite having a strong effect on URX. Conversely, the allele with the most penetrant BAG defects (*hmn4*) exhibits the least penetrant URX defects, albeit still quite severe. These results suggest that, although they require the same factors for dendrite extension, URX and BAG are likely to exhibit some developmental differences.

Importantly, a *grdn-1* deletion allele (*tm6493*) causes completely penetrant embryonic lethality (Nechipurenko et al., 2016), suggesting this gene plays an essential role in early development. This leads to three inferences. First, our genetic approach has revealed a role in dendrite development that would have been precluded from study using a null allele. Second, because our mutants are viable (*ns303*, <2% embryonic lethality, n=138), they presumably retain some *grdn-1* activity (i.e., they are hypomorphic alleles). Third, these alleles might therefore provide clues as to which isoforms or protein regions are specifically required for dendrite extension and not for other aspects of early development. For example, all of our alleles disrupt the GRDN-1a isoform while some of them (*ns303, ns302, hmn7, hmn8)* spare the GRDN-1b or GRDN-1c isoforms. This suggests that GRDN-1a is likely the most relevant for dendrite extension and that neither GRDN-1b nor GRDN-1c is sufficient. Consistent with this prediction, we found that GRDN-1a cDNA can rescue URX dendrite extension in the *grdn-1(ns303)* background (Fig. S4). Interestingly, one allele that strongly affects URX – but not BAG – dendrite extension (*hmn8*) is predicted to truncate GRDN-1 just 26 aa from its carboxy-terminus. This short region includes a PDZ-domain-binding motif (Gly-Cys-Val), suggesting that this motif is required for URX dendrite extension but not other aspects of early development. Consistent with this prediction, deleting this motif from the GRDN-1a cDNA abrogates its ability to rescue URX dendrite extension defects (Fig. S4). We chose to focus on the *ns303* allele, as it is predicted to truncate GRDN-1a close to its amino terminus (after the first seven codons) and causes highly penetrant defects in both URX and BAG without embryonic lethality.

In summary, it is intriguing that our screen exclusively identified alleles of *sax-7* and *grdn-1*, and that similar phenotypes are not caused by mutations in known factors that interact with SAX-7 and GRDN-1 during neuronal development. However, in addition to their roles in neurons, SAX-7 and GRDN-1 are also expressed broadly in epithelial tissues during embryonic development (Chen et al., 2001; Nechipurenko et al., 2016). An interesting possibility is that factors that interact with them during epithelial development might promote URX and BAG dendrite extension but were not identified in our screens due to other, essential roles.

### SAX-7 and GRDN-1 are not broadly required for dendrite extension

As SAX-7 and GRDN-1 both act broadly in development, we sought to determine if their function in dendrite extension is specific to URX and BAG. We therefore asked whether *sax-7* or *grdn-1* affect the morphogenesis of other sensory neurons that extend unbranched dendrites to the nose.

First, we examined a panel of strains bearing cell-type-specific fluorescent markers for three of the 12 amphid neurons as well as three other classes of sensory neurons that extend dendrites through glial pores (CEP, IL, OL). We found that most of these neurons are unaffected by *sax-7* and *grdn-1* (Fig. 4B,D). However, the amphid neurons AWA and AWB exhibited weakly-penetrant dendrite extension defects in *grdn-1* (Figs. 4, S5), consistent with a previous report that noted aberrant branching and other morphological defects (Nechipurenko et al., 2016). These defects could reflect a primary role for GRDN-1 during AWA and AWB dendrite extension, or could be secondary to defects in basal body positioning in these neurons, as many mutants that disrupt the ciliary basal body or transition zone also produce some amphid dendrite defects (CHE-10, Fig. S3B; MKS-1, NPHP-4 (Williams et al., 2008); CCEP-290, MKSR-2, NPHP-4 (Schouteden et al., 2015); MKS-5, MKS-6, NPHP-1, NPHP-4 (Williams et al., 2011)). Importantly, we found that AWA and AWB dendrite lengths are not affected by *sax-7*, and another amphid neuron (AWC) exhibits no detectable dendrite extension defects in either mutant (Fig. S5).

**Fig. 4.**
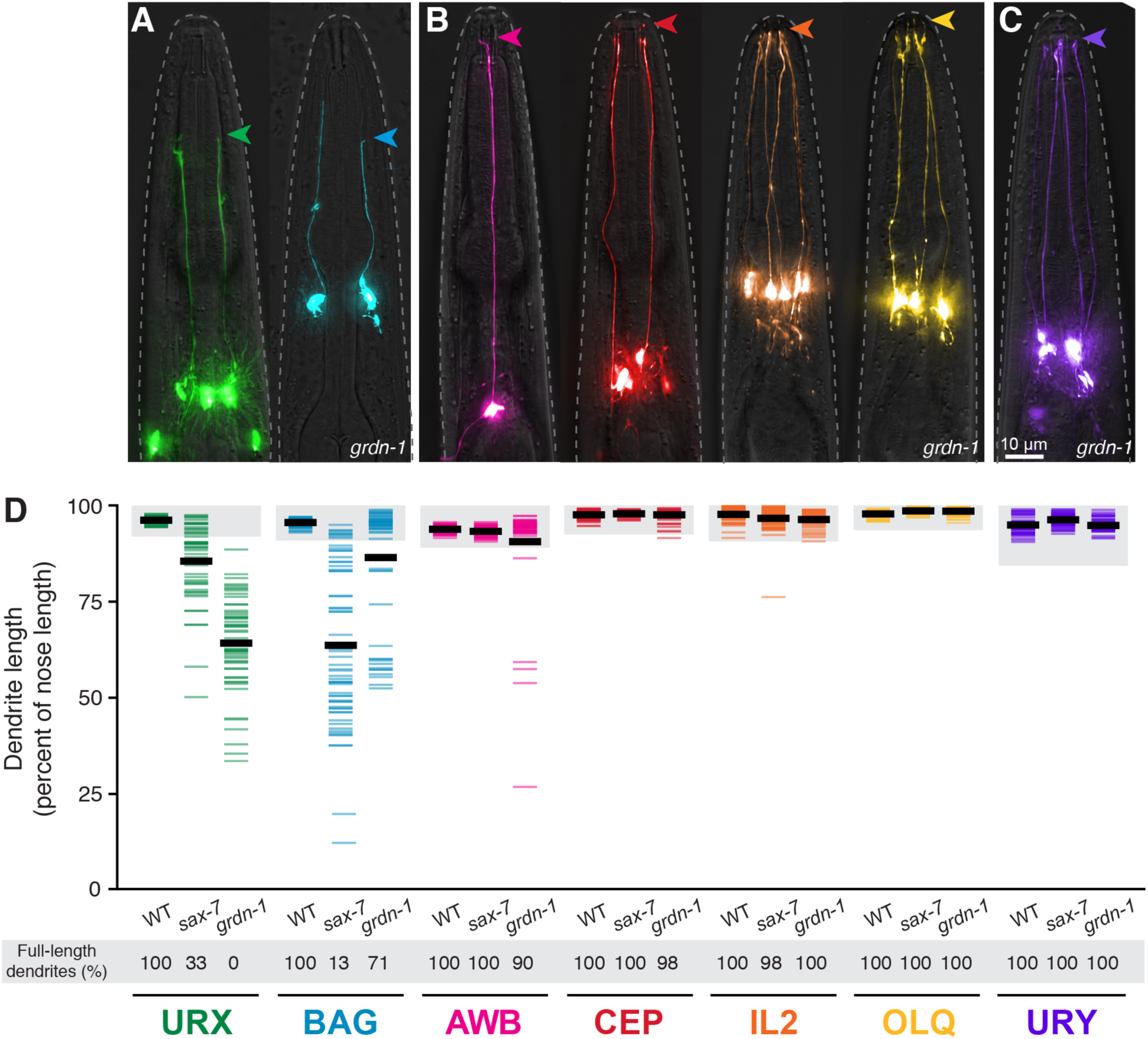
*sax-7* and *grdn-1* preferentially affect URX and BAG dendrite extension. (A, B) *grdn-1* animals expressing fluorescent markers for (A) neurons that make specialized wrapping contacts with the ILso glia: URX (green, *flp-8*pro) and BAG (blue, *flp-17*pro); (B) neurons that enter glial pores: amphid (AWB, pink, *str-1*pro; two additional amphid neurons, AWA and AWC, are in Fig. S5), CEP (red, *dat-1*pro), IL2 (orange, *klp-6*pro), OLQ (yellow, *ocr-4*pro); and (C) a neuron that does neither: URY (purple, *tol-1*pro). Arrowheads, dendrite endings. (D) Quantification of dendrite lengths in the indicated genotypes, expressed as a percentage of the distance from cell body to nose. Colored bars represent individual dendrites (n≥47 per genotype); black bars represent population averages. Shaded region represents wild-type mean ± 5 s.d. for each neuron type and the percentage of dendrites in this range (“full length dendrites”) is indicated below the plots.

We also examined URY, another neuron that extends an unbranched dendrite to the nose. Like URX and BAG, the URY dendrite ending interacts closely with some glia (although not the ILso glia) but does not enter a glial pore, is not exposed to the external environment, and does not require DYF-7 (Ward et al., 1975; Doroquez et al., 2014; Low et al., 2019). We found that URY dendrite extension is not affected by either *sax-7* or *grdn-1* (Fig. 4C, D).

Thus, despite the widespread roles of *sax-7* and *grdn-1* in developmental patterning, their effects on dendrite extension appear relatively restricted to URX and BAG, and might reflect mechanisms specific to how these neurons form their dendrites.

### SAX-7 and GRDN-1 promote dendrite growth during embryo elongation

To determine how URX and BAG form their dendrites – and, in particular, whether they develop by retrograde extension – we visualized dendrite development in wild-type and mutant embryos using a fluorescent reporter expressed in these neurons shortly after their birth (*egl-13*pro:GFP) (Fig. 5) (Gramstrup Petersen et al., 2013). URX and BAG develop at a stage when the embryo rapidly twitches, making imaging difficult. We employed two approaches to circumvent this problem. First, we used dual-view inverted selective-plane illumination microscopy (diSPIM) (Wu et al., 2011; Kumar et al., 2014), which allows optical volumes to be acquired faster than the embryo twitches, to perform time-lapse imaging of a small number of embryos (Fig. S6, Movies S1 and S2). However, due to the variability of the mutant phenotype across individuals, we also wanted to use a method that would allow sampling of larger populations. We therefore took advantage of our previous anecdotal observations that ultraviolet light causes rapid embryonic arrest (see Methods), and used this approach to arrest and image >70 wild-type or mutant embryos at various developmental stages. We then arranged these embryos into a pseudo-time-course based on the highly stereotyped developmental features of *C. elegans* embryogenesis (Fig. 5A-C).

**Fig. 5.**
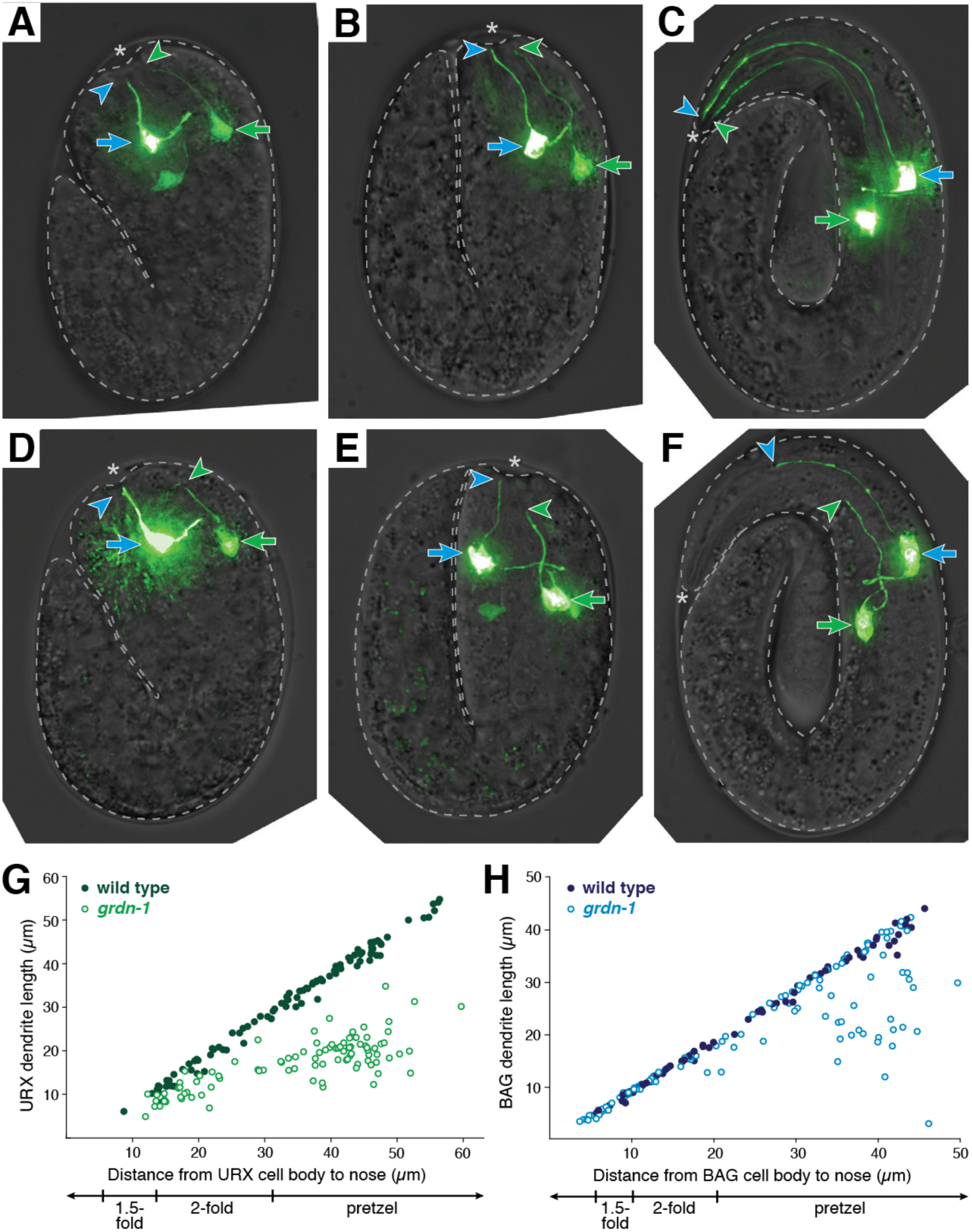
URX and BAG dendrite length defects arise during embryo elongation. (A-F) URX and BAG dendrite extension in (A-C) wild-type and (D-F) *grdn-1* embryos at (A, D) 1.5-fold, (B, E) 2-fold, and (C, F) pretzel stages. Embryonic URX and BAG were visualized using *egl-13*pro:GFP and distinguished by their cell body positions (URX cell body is more posterior, green arrow; BAG cell body, blue arrow). URX and BAG dendrite endings are marked with green and blue arrowheads, respectively. Asterisk, embryo nose. (G, H) Dendrite lengths plotted as a function of the distance from cell body to nose for (G) URX and (H) BAG in wild-type (filled circles) and *grdn-1* (open circles) embryos. n≥70 per genotype.

In wild-type embryos, at the earliest stages where we can conclusively identify URX and BAG, the dendrites of both neurons are partially extended and contact the presumptive nose (Fig. 5A). At this stage, neither cell has a migratory appearance – for example, they lack the lamellipod-like structure seen in developing amphid neurons (Heiman and Shaham, 2009), and BAG has already extended a short medial axon. Interestingly, the dendrite endings do not converge, suggesting that their shared attachments to the lateral ILso glial cell observed in the mature structure may not have developed yet. Next, as embryo elongation begins, the dendrites appear to stretch as the distance from the cell bodies to the nose increases (Fig. 5B). We found that dendrites increase at least five-fold in length during this period, from ∼5-10 µm to ∼50 µm, exactly keeping pace with the increasing distance from the cell bodies to the nose (Fig. 5C, G, H). Thus, URX and BAG develop using retrograde extension, with the nascent dendrites attaching near the presumptive nose and then growing by stretch as the cell bodies move farther away during embryo elongation.

To determine if *sax-7* and *grdn-1* are required for this form of retrograde extension, we examined the development of mutant embryos. We focused on *grdn-1* for analysis, because the URX phenotype is fully penetrant, but we also observed similar results with *sax-7* (Fig. S6). In *grdn-1* embryos, we found that URX and BAG initially adopt their normal polarized morphology and form nascent dendrites that contact the presumptive nose (Fig. 5D). However, during embryo elongation, the dendrite endings become displaced from the nose and fail to extend normally (Fig. 5E, F). URX dendrites remained in contact with the nose up to a length of ∼15-20 µm; however, as embryo elongation proceeded, they failed to maintain contact with the nose and were always shorter than wild type (Fig. 5G). BAG dendrites exhibited less penetrant defects, consistent with the lower penetrance observed in the mature structure. Similar to URX, dendrite extension defects in BAG were almost never seen at earlier stages (cell body <20 µm from the nose) but, as embryo elongation proceeded, ∼30% of dendrites became displaced from the nose (Fig. 5H). Thus, URX and BAG dendrite endings anchor to the presumptive nose using SAX-7 and GRDN-1, a mechanism of retrograde extension that is molecularly distinct from that employed by amphid neurons. An intriguing possibility is that contacts with glia are responsible for dendrite tip anchoring; however, because we do not have tools to visualize ILso glia in the embryo, we cannot directly test this idea. It is also possible that URX and BAG dendrites attach to other partners during development and then attain their specialized glial contacts later.

Finally, we asked how *sax-7* and *grdn-1* affect the post-embryonic scaling of dendrite length. Within ∼48 h of hatching, the head increases two-fold in length as animals grow from the first (L1) to fourth (L4) larval stage. We found that mutant dendrites generally scaled appropriately to keep pace with larval growth over this period, despite their overall shorter lengths (Fig. S7A). We also noted that URX and BAG dendrite lengths were uncorrelated, suggesting that defects in URX are not secondary consequences of defects in BAG or vice versa (Fig. S7B). Together, our results imply that *sax-7* and *grdn-1* specifically disrupt the retrograde extension of URX and BAG dendrites during embryo elongation.

### SAX-7 can act both in neurons and glia to promote URX and BAG dendrite extension

Because vertebrate homologs of SAX-7 can mediate neuron-glia adhesion, we reasoned that SAX-7 might also mediate neuron-glia adhesion to anchor dendrites at the developing nose. To test this idea, we expressed SAX-7 cDNA under neuron- or glia-specific promoters in a *sax-7* mutant background and quantified the number of full-length URX and BAG dendrites. As a positive control, we observed nearly complete rescue when SAX-7 cDNA was placed under control of a broadly-expressed embryonic promoter (Fig. 6A, B, *grdn-1*pro). We found that expression using URX-, BAG- or glia-specific promoters (*flp-8*pro, URX; *flp-17*pro, BAG; *itx-1*pro, glia) (Kim and Li, 2004; Haklai-Topper et al., 2011) produced moderate rescue, whereas combined expression in URX and glia, or BAG and glia, produced much more efficient rescue (neurons with full-length dendrites: 92%, URX; 78%, BAG) (Fig. 6A,B). The lack of complete rescue could reflect additional roles for SAX-7 in other cell types, or issues related to the developmental timing of the cell-specific promoters we used. These results suggest that SAX-7 promotes dendrite anchoring by acting both in the URX and BAG neurons as well as in glia.

**Fig. 6.**
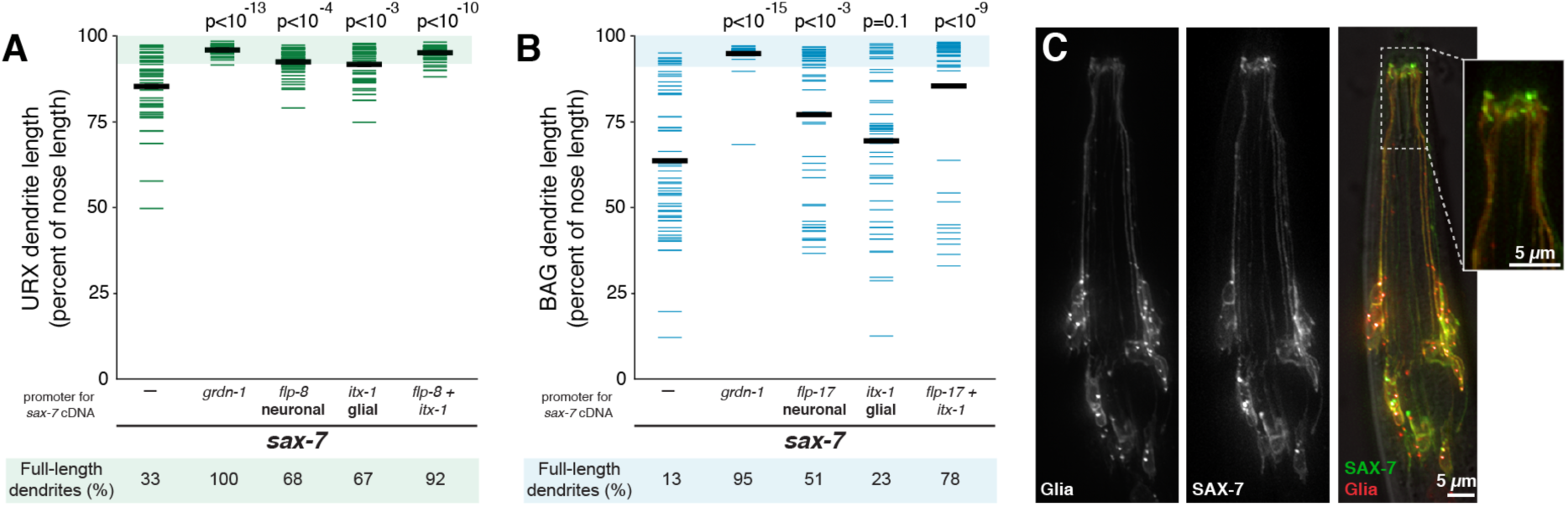
SAX-7 can act both in neurons and glia to promote dendrite extension. (A, B) Transgenes containing SAX-7S cDNA under the control of promoters expressed in neurons, glia, both together, or no transgene (–), were introduced into *sax-7* animals. (A) URX and (B) BAG dendrite lengths were measured in each strain as a percentage of the distance from cell body to nose. Colored bars represent individual dendrites (n≥49 per genotype); black bars represent population averages. p-values (Wilcoxon Rank Sum test) compared to *sax-7* with no transgene (–) are shown at top. Rescue was greater when SAX-7S cDNA was expressed in neurons and glia together compared to neurons only (URX, p<10^-5^; BAG, p<10^-3^) or glia only (URX, p<10^-4^; BAG, p<10^-5^). Shaded region represents wild-type mean ± 5 s.d. for each neuron type and the percentage of dendrites in this range (“full length dendrites”) is indicated below the plots. (C) Single-wavelength and merged images of animals expressing SAX-7S-YFP and the membrane marker myristyl-mCherry under control of a glia-specific promoter (*itx-1*pro). Inset: SAX-7S-YFP is sometimes enriched in glial endings, where the glia contact URX and BAG.

Next, to ask whether SAX-7 localizes to specific regions within glia, we expressed a functional YFP-tagged SAX-7 cDNA under control of a glial-specific promoter (*itx-1*pro), together with myristyl-mCherry (myr-mCherry) as a reference (Fig. 6C). We found that SAX-7 localizes across the glial surface and often appears enriched at sites near the nose (Fig. 6C). This enrichment was not seen in all individuals, but is consistent with the possibility that SAX-7 could directly mediate dendrite-glia attachment.

### GRDN-1 can act non-autonomously in glia to promote dendrite extension

Next, we performed a similar analysis of GRDN-1. In contrast to the above results, we observed almost no rescue, or even mildly enhanced defects, when the GRDN-1a cDNA was placed under control of URX- or BAG-specific promoters (Fig. 7A, S8A). This suggests that GRDN-1, unlike SAX-7, does not act in the neurons themselves to promote dendrite extension. We recapitulated this result using two other neuronal promoters (*egl-13*pro, expressed in URX and BAG shortly after the cells are born; *rab-3*pro, pan-neuronal) (Fig. 7A, S8A) (Nonet et al., 1997; Gramstrup Petersen et al., 2013).

**Fig. 7.**
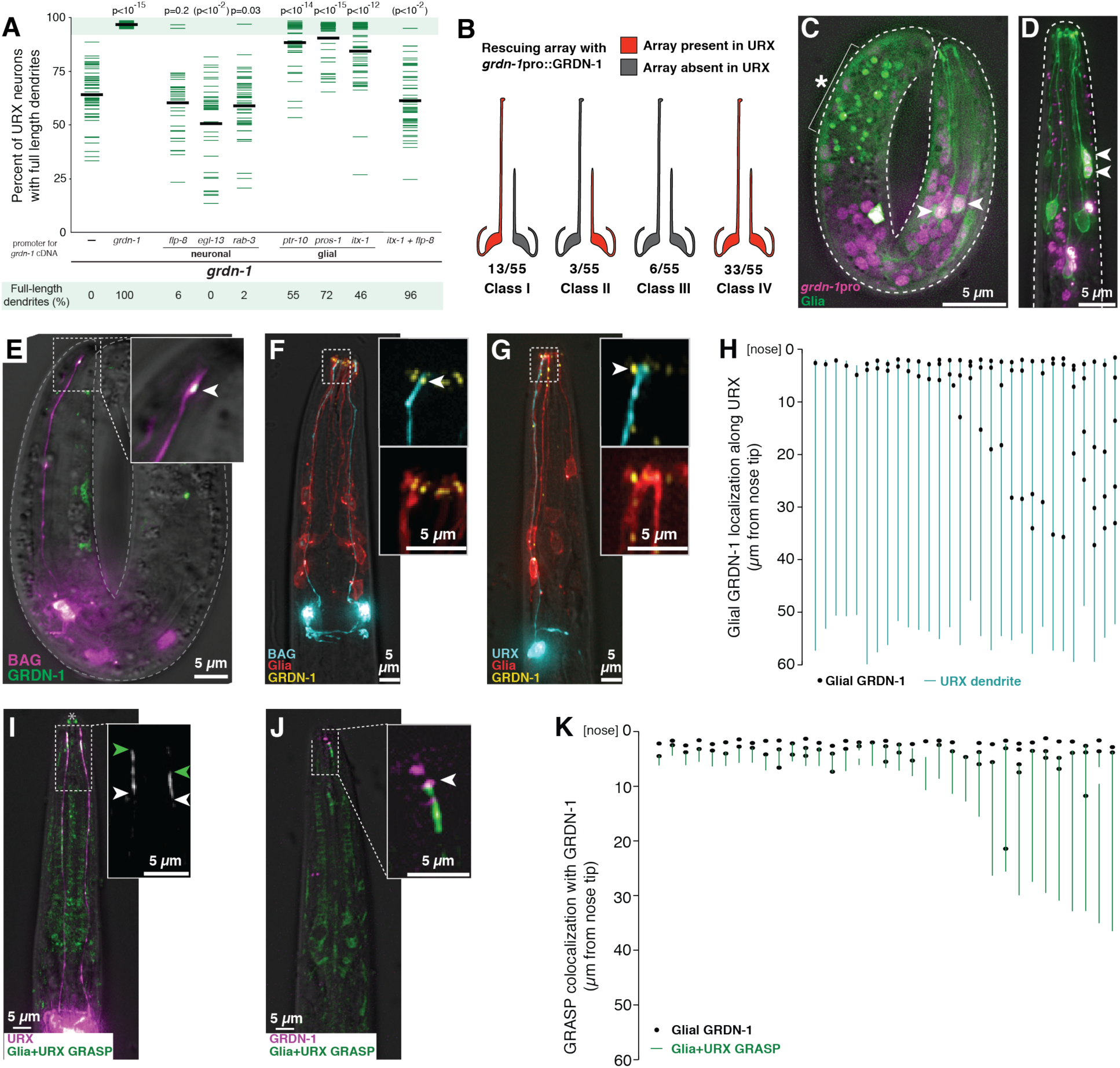
GRDN-1 can act in glia and localizes in puncta at dendrite contacts. **(A)** Transgenes containing GRDN-1a cDNA under control of the indicated promoters, or no transgene (–), were introduced into *grdn-1* animals and URX dendrite lengths were measured as a percentage of the distance from the cell body to the nose. p-values (Wilcoxon Rank Sum test) compared to *grdn-1* with no transgene (–) are shown at top. Parentheses indicate that, for unknown reasons, expression using *egl-13*pro enhances the defects. Colored bars represent individual dendrites (n≥34 per genotype); black bars represent population averages. Shaded region represents wild-type mean ± 5 s.d. and the percentage of dendrites in this range (“full length dendrites”) is indicated below the plots. Data for BAG is in Fig. S8. (B) Mosaic analysis was performed using *grdn-1* animals bearing a stably-integrated URX marker (*flp-8*pro:GFP) to assess dendrite length and an extrachromosomal transgene, which is subject to stochastic loss during cell division, containing *grdn-1*pro:GRDN-1a and the URX marker *flp-8*pro:CFP to assess the ability of GRDN-1 to act cell-autonomously. Animals in which only the left or right URX dendrite was full-length were selected as probable mosaics, and were scored for the presence (red) or absence (gray) of the rescuing array in each URX neuron. (C, D) Wild-type embryo (C) and L1 stage larva (D) expressing a histone-mCherry fusion protein under control of the *grdn-1* promoter (*grdn-1*pro:*his-24-mCherry*, magenta) and a membrane-localized GFP to mark glia (*itx-1*pro:myristyl-GFP, green). Arrowheads, examples of cells expressing both markers. Asterisk and bracket, background auto-fluorescence due to gut granules. (E-H) The localization of glial-expressed GRDN-1a relative to URX and BAG was visualized (E) in embryos, by expressing sfGFP-GRDN-1a in glia (*itx-1*pro, green) together with an embryonic BAG marker (*egl-13*pro:mCherry, purple); and (F, G) in L1 stage larvae, by expressing YFP-GRDN-1a (*itx-1*pro, yellow) and myristyl-mCherry (*itx-1*pro, red) in glia together with CFP (blue) in BAG (F, *flp-17*pro) or URX (G, *flp-8*pro). Arrowheads, glial-expressed puncta of GRDN-1a near dendrite endings. (H) The positions of glial-expressed GRDN-1a puncta were quantified relative to the URX dendrite. Each line represents a single URX dendrite, and adjacent YFP-GRDN-1a puncta are shown as black dots. n=30. (I-K) GRDN-1a puncta were visualized relative to the dendrite-glia contact. (I) Regions of direct dendrite-glia contact were visualized using a GFP-reconstitution assay (GRASP) targeting URX and glia (*itx-1*pro:CD4-spGFP1-10 and *flp-8*pro:CD4-spGFP11, green). URX is also marked by mCherry (*flp-8*pro, magenta). Arrowheads mark the anterior and posterior extents of the GRASP signal. Due to the faint signal, prominent autofluorescence at mouth (asterisk) and elsewhere in the head is visible. (J) The localization of glial-expressed GRDN-1a relative to the URX-glia contact was visualized by expressing mApple-GRDN-1a in glia (*itx-1*pro, magenta) together with GRASP (green). Arrowhead, glial-expressed punctum of GRDN-1a at the anterior extent of the dendrite-glia contact revealed by GRASP. (K) The positions of glial-expressed GRDN-1a puncta were quantified relative to the dendrite-glia contact. Each line represents one URX-glia pair as revealed by GRASP, and adjacent mApple-GRDN-1a puncta are shown as black dots. Multiple dots may reflect expression of the promoter in several (∼6-12) glial cells. n=35.

By comparison, we observed moderate rescue of URX dendrite extension using any of three glial promoters (neurons with full-length dendrites: 55%, *ptr-10*pro; 72%, *pros-1*pro; 46%, *itx-1*pro) (Fig. 7A) (Yoshimura et al., 2008; Haklai-Topper et al., 2011; Murray et al., 2012). As discussed above, incomplete rescue could be due to inappropriate timing of expression or a requirement for *grdn-1* in additional cells. These results suggest that GRDN-1 is able to act non-autonomously in glia to promote URX dendrite anchoring.

Because *grdn-1* causes more weakly penetrant defects in BAG, we could not determine its site of action for BAG dendrite anchoring. However, we note that some glial promoters resulted in partial rescue of BAG comparable to that seen for URX – for example, *ptr-10*pro:GRDN-1 reduced the fraction of affected dendrites by about half in both cases (BAG, 16% affected compared to 29% in controls, Fig. S8A; URX, 45% affected compared to 100% in controls, Fig. 7A). For reasons we do not understand, combined expression of GRDN-1 in neurons and glia abrogated the rescue, or was even slightly detrimental, for both URX and BAG dendrite extension (Fig. 7A, S8A).

To further test whether GRDN-1 acts non-autonomously, we used mosaic analysis to interrogate the activity of *grdn-1* expressed under control of its own promoter (Fig. 7B). Briefly, we generated *grdn-1* animals bearing an unstable extrachromosomal transgene consisting of *grdn-1*pro:GRDN-1a cDNA and a fluorescent reporter (*flp-8*pro:CFP) used to visualize the presence of this transgene in URX. To identify rare mosaic animals in which the transgene has undergone stochastic loss in certain cell lineages, we isolated individuals in which the left- and right-hand URX neurons are of different lengths (i.e. one short dendrite and one full-length dendrite). We reasoned that, if *grdn-1* acts cell-autonomously, then the URX neuron with a full-length dendrite would always carry the transgene (*grdn-1(+),* Fig. 7B, red), while the neuron with a short dendrite would almost always lack the transgene (*grdn-1(–)*, Fig. 7B, gray). However, in addition to this class of animals (Class I, 23%), we also observed two classes of animals in which URX neurons had full-length dendrites but lacked the transgene (Classes II and III, 16% together), indicating that *grdn-1* is not necessary in URX for rescue (Fig. 7B). Similarly, we observed two classes of animals in which URX neurons had short dendrites despite carrying the transgene (Classes II and IV, 65% together), indicating that *grdn-1* is not sufficient in URX for rescue (Fig. 7B). These results support the conclusion that *grdn-1* promotes URX dendrite extension in a non-autonomous manner.

Finally, we asked if *grdn-1* is normally expressed in glia. We generated a *grdn-1* transcriptional reporter and found that it is broadly expressed in the embryo from late gastrulation onwards. By co-expressing a glial marker in embryos (*itx-1*pro(1 kb), see Methods), we found that this expression includes a subset of glia (Fig. 7C). This expression can be detected in newly-hatched animals (Fig. 7D) but is absent in older larvae and adults. Importantly, *grdn-1* does not grossly disrupt the specification or morphogenesis of these glia (Fig. S2B), although the protrusions at the glial ending appeared to be diminished or absent when URX and BAG dendrites were both short (Fig. S8C).

Taken together, our results suggest that *grdn-1* is expressed broadly at the time of dendrite extension, including in glia, and that its expression in glia is sufficient to non-autonomously promote dendrite extension.

### GRDN-1 localizes in glia to puncta near dendrite contacts

To examine GRDN-1 localization, we generated a functional GFP-tagged GRDN-1a cDNA. Consistent with previous results, we observed numerous puncta at the nose when this construct was expressed under its own promoter (Fig. S8B). These puncta were previously shown to localize to the basal bodies at the ciliated endings of sensory dendrites (Nechipurenko et al., 2016). However, as glia are not ciliated, we were curious how GRDN-1 would localize in these cells. We therefore expressed GFP-GRDN-1 under the control of a glial-specific promoter. In embryos and newly-hatched larvae, we found that glial GRDN-1 localizes in puncta near URX and BAG dendrite endings (Fig. 7E-G). This localization did not depend on *sax-7* (Fig. S8D). Quantifying the positions of puncta relative to the URX dendrite revealed that the dendrite always terminates near a punctum of glial GRDN-1 (n=30/30, Fig 7H).

Because these puncta were not bright enough for the structured illumination approach we used previously, we developed an alternative strategy to test if they represent dendrite-glia contacts. Briefly, we adapted the GRASP system in which complementary fragments of GFP are expressed on the surfaces of neighboring cells, such that fluorescence is reconstituted only where the cells come into molecular contact (Feinberg et al., 2008). We expressed the GRASP fragments in URX and glia (URX+Glia GRASP) together with a reference marker in URX. We observed GRASP signal ranging in length from ∼3 to 30 µm along the distal dendrite (Fig. 7I), consistent with our structured illumination images showing the distal portion of URX tightly bundled with the ILso glia. As a control, no signal was observed when either GFP fragment was expressed alone. By expressing this GRASP reporter together with glial-specific mApple-GRDN-1, we found that the GRDN-1 puncta almost always mark the anterior extent of the dendrite-glia contact (n=34 of 35 animals; Fig. 7J, K). Thus, distinct from its role at basal bodies in ciliated neurons, GRDN-1 localizes in glia to puncta positioned at dendrite-glia contacts.

## DISCUSSION

### A novel molecular mechanism for retrograde extension

Here, we establish a novel genetic model for dissecting dendrite-glia interactions at the single cell level. We find that the mature URX and BAG dendrites form specialized membranous contacts with a single glial cell, the lateral ILso. During development, these dendrites grow by retrograde extension, attaching near the nose and then stretching out during embryo elongation. Dendrite anchoring requires the neuron-glia adhesion molecule SAX-7, as well as GRDN-1, which can act non-autonomously in glia and localizes in puncta near dendrite-glia contacts. In the absence of these factors, dendrites detach from the nose and do not properly extend during embryo elongation. Thus, this study identifies conserved factors that may promote dendrite-glia adhesion in order to drive dendrite growth by retrograde extension.

Interestingly, this represents a novel molecular mechanism for retrograde extension. While URX and BAG superficially resemble the amphid and other sensory neurons – they extend unbranched dendrites to the nose where they interact with glia – they exhibit highly divergent features and develop using distinct cellular and molecular mechanisms (Fig. 8). Amphid dendrites develop as part of an epithelium: they are exposed to the environment, form epithelial tight junctions with neighboring glia, exhibit apical-basal polarity, and develop using apical ECM factors that resemble those found in other epithelia (Low et al., 2019). In contrast, URX and BAG are not exposed to the environment, do not form tight junctions, and do not require the apical ECM factors employed by the amphid (Ward et al., 1975; Doroquez et al., 2014). URX and BAG dendrites instead attach to the basolateral surface of ILso glia and develop using SAX-7 and GRDN-1, conserved molecules that promote cell adhesion in the mammalian brain. Interestingly, retrograde extension has recently been observed in the developing zebrafish nervous system (Breau et al., 2017), suggesting that this strategy for neurite growth may be more widespread than was previously appreciated. It will be important to determine if SAX-7/GRDN-1-dependent mechanisms also mediate retrograde extension in vertebrates, or if there are yet other molecular mechanisms for retrograde extension.

**Fig. 8.**
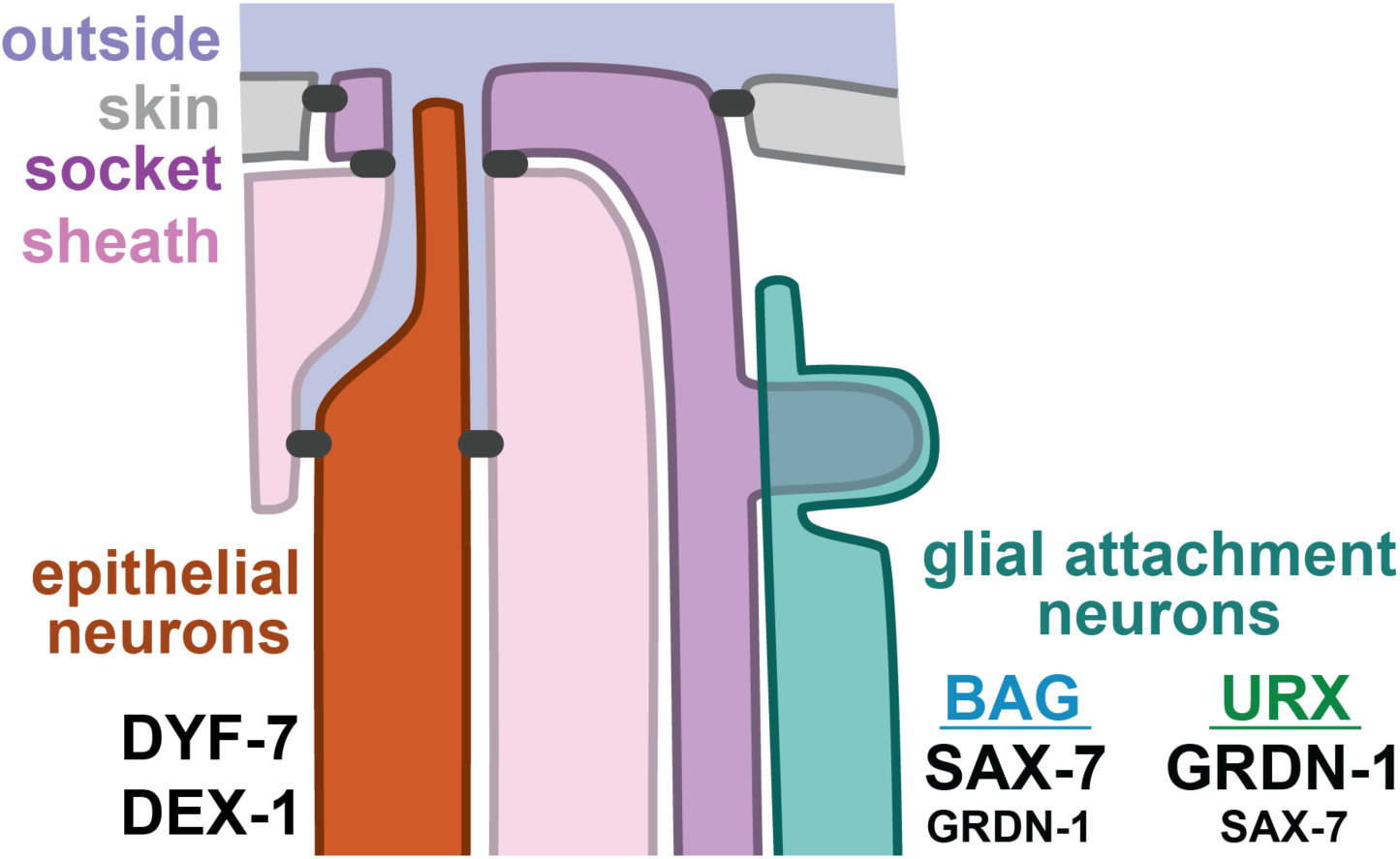
Distinct molecular mechanisms of retrograde extension. Different classes of head sensory neurons undergo retrograde extension using distinct cellular and molecular mechanisms. Epithelial neurons (24 amphid, 12 IL, six OL, four CEP) exhibit tight junctions (black) that separate an outward- or lumenal-facing apical surface from a basal surface that is exposed to the pseudocoelom, and undergo retrograde extension through mechanisms that resemble the morphogenesis of narrow epithelial tubes, including the use of apical extracellular matrix proteins (DYF-7 and DEX-1). In contrast, glial attachment neurons (two BAG, two URX) undergo retrograde extension using a distinct molecular mechanism in which SAX-7 and GRDN-1 anchor dendrite endings at the developing nose. BAG exhibits a greater dependence on SAX-7, while URX exhibits a greater dependence on GRDN-1. Dendrite extension in the other unbranched sensory neurons of the head (four URA, two URB, four URY) remains to be explored.

Interestingly, the URX dendrite is not restricted to growing only by retrograde extension. We previously identified a separate class of dendrite overgrowth mutants in which the URX dendrite forms normally during embryogenesis but then gets longer as the animal ages, ultimately wrapping around the nose and extending posteriorly along the head, up to 150% of its wild-type length (McLachlan et al., 2018). These results suggest a possible developmental switch from retrograde extension in the embryo to tip-directed anterograde extension in larvae and adults. In the developing zebrafish nervous system, axons that initially grow by retrograde extension later undergo a second phase of anterograde growth, during which they grow away from the surface of the brain and towards their targets within it (Breau et al., 2017). This suggests that a developmental switch between retrograde and anterograde growth may also be conserved across species.

The model of dendrite-glia interaction we have established may also provide insight into other aspects of mammalian brain development. L1CAM/SAX-7 and CCDC88C/GRDN-1 are two of the genes most frequently disrupted in congenital hydrocephalus, a rare developmental brain disorder (Drielsma et al., 2012; Adle-Biassette et al., 2013; Nahorski et al., 2016). Could the molecular mechanisms for congenital hydrocephalus be related to the overlapping roles of SAX-7 and GRDN-1 described here? Congenital hydrocephalus is characterized by enlargement of the brain ventricles, which arise developmentally from the lumen of the embryonic neural tube. Loss of CCDC88C leads to apical constriction defects in the neuroepithelial cells that line the lumen of the developing neural tube, and planar cell polarity (PCP) defects in the epithelial radial glial cells that subsequently line the surfaces of newly-formed ventricles (Takagishi et al., 2017; Marivin and Garcia-Marcos, 2019; Marivin et al., 2019). These defects are not obviously related to the phenotypes described here. However, it is intriguing to note that ILso is an epithelial glial cell. It is therefore possible that it shares some underlying developmental features with vertebrate neuroepithelial cells or epithelial radial glia.

### Possible molecular mechanisms for SAX-7 and GRDN-1

By what molecular mechanism do SAX-7 and GRDN-1 promote dendrite extension? A simple hypothesis would be that they form a protein complex that attaches URX and BAG dendrites to the lateral ILso glial cell. However, several lines of evidence suggest that the mechanism is more complicated. First, SAX-7 and GRDN-1 do not have domains known to interact with each other and, in contrast to what would be expected for two members of a protein complex, they do not exhibit the same localization pattern in glia. Second, loss of SAX-7 is not phenotypically equivalent to loss of GRDN-1 – for example, SAX-7 more strongly affects BAG while GRDN-1 more strongly affects URX – which likely would not be the case if these genes acted in a single complex. Third, embryonic BAG and URX dendrites do not converge to a single site at the nose, as would be expected if they were both already attached to the lateral ILso glial cell, suggesting the dendrites might transiently attach to different cellular partners during development before attaining their mature structure. These observations suggest that SAX-7 and GRDN-1 may act in parallel, rather than in a single complex, to promote dendrite extension.

A possible clue to their mechanism of action may come from studies of epithelial development, in which each of these proteins has been found to act upstream of epithelial cell junctions. SAX-7 is expressed broadly in epithelia, localizes to epithelial cell junctions, and can act redundantly with the classical epithelial cell junction protein HMR-1/cadherin during early development (Chen et al., 2001; Grana et al., 2010). GRDN-1 is partly required for localization of the epithelial cell junction marker AJM-1 and, in *Drosophila,* Girdin localizes to epithelial adherens junctions, physically interacts with the cadherin-catenin complex protein α-catenin, and is required to maintain epithelial cell adhesion during large-scale morphogenetic movements (Ha et al., 2015; Houssin et al., 2015; Nechipurenko et al., 2016).

These observations suggest two possibilities. First, URX and BAG dendrites might attach to glia through modified epithelial junctions, for example using the same cell junction machinery as epithelia but in an attenuated form that is more susceptible to disruption. This is an attractive idea because neurons and glia derive developmentally and evolutionarily from epithelia and many glia retain epithelial characteristics (including ILso, as well as radial glia and Müller glia in vertebrates). Alternatively, URX and BAG dendrites might develop using bona fide epithelial interactions that are lost as the structure matures. Recent studies have shown that other *C. elegans* neurons engage in transient cellular rosettes, in which cells polarize towards a central vertex and exhibit epithelial characteristics before resolving into their mature arrangement (Fan et al., 2019). It will therefore be paramount to identify the early cell-cell interactions that shape URX and BAG, and to understand how they give rise to the specialized dendrite-glia contacts observed in the mature structure.

## ACKNOWLEDGMENTS

We thank Valeri Thomson for assistance with the genetic screen that isolated *ns302* and *ns303*, and Shai Shaham for generously providing laboratory resources, advice, and supervision in support of the isolation and initial characterization of these mutants; Anique Olivier-Mason, Inna Nechipurenko and Piali Sengupta for sharing reagents, unpublished information, and comments on the manuscript; John Murray for advice on promoter expression patterns; Ikue Mori for *sax-7* cDNA; Cori Bargmann for GRASP constructs; Chris Li for advice and reagents related to *flp-8* and *flp-17*; Alex Boyanov and Oliver Hobert for whole-genome sequencing; Talley Lambert, Anna Payne-Tobin Jost, and Jennifer Waters (Harvard Medical School Cell Biology Microscopy Facility) for their assistance with diSPIM and SIM acquisition; Lin Shao (Yale University) for CUDA-accelerated 3D-SIM reconstruction code; and Lisa Goodrich and members of the Heiman laboratory for comments on the manuscript. Some strains were provided by the CGC, which is funded by NIH Office of Research Infrastructure Programs (P40 OD010440). We are grateful for funding from the National Science Foundation Graduate Research Fellowship Program (I.G.M. and E.R.C.), NIH F31NS103371 (E.R.C.), NIH R01GM108754 (M.G.H.), NIH R01NS112343 (M.G.H.), and a March of Dimes Basil O’Connor Starter Scholar Research Award (M.G.H.).

## MATERIALS AND METHODS

### Strains and Plasmids

All *C. elegans* strains were constructed in the N2 background and cultured at 20°C on nematode growth media (NGM) plates seeded with *E. coli* OP50 bacteria (Brenner, 1974). Transgenic strains were generated with standard techniques (Mello and Fire, 1995) with injection of 100 ng/µL of DNA (5-50µL per plasmid). Information for all strains used in this study is listed in Table S1, plasmids are listed in Table S2, and primers of general interest are listed in Table S3.

### Generation of glial expression constructs

To visualize inner and outer labial glial cells, we amplified a ∼2.9 kb promoter fragment upstream of *itx-1* from N2 genomic DNA and used it to drive the expression of GFP. Consistent with published results (Haklai-Topper et al., 2011), this promoter drives expression in glial cells and throughout the gut. As the strong gut expression obscured signal from the head in embryos, we identified a 1,031 bp region beginning at the 5’ end of the ∼2.9 kb *itx-1* promoter that drives expression in glia, but not in the gut, when placed upstream of a *myo-2* minimal promoter (Okkema et al., 1993). See Table S3 for primer sequences.

To visualize inner labial socket glial cells specifically, we used a promoter fragment from the *grl-18* gene that was reported to be expressed in a combination of IL or OL socket glia (Hao et al., 2006). We amplified a ∼3 kb promoter fragment upstream of *grl-18* from N2 genomic DNA and used it to drive expression of YFP. To confirm expression in IL socket glia, we co-expressed *grl-18*pro:YFP with the BAG reporter *flp-17*pro:CFP and the IL2 neuron reporter *klp-6*pro:mCherry. We found that the IL2 neurons protrude through the *grl-18+* cells and that BAG neurons associate with the YFP-labeled cells, thus demonstrating that the *grl-18* promoter drives expression specifically in IL socket glia.

### Forward genetic screens

We isolated alleles of *grdn-1* and *sax-7* in three independent, nonclonal, visual genetic screens. In all screens, L4 stage animals were mutagenized using 70 mM ethyl methanesulfonate (EMS, Sigma) at approximately 22°C for 4 hours. Nonclonal F2 progeny were examined on a Nikon SMZ1500 stereomicroscope with an HR Plan Apo 1.6x objective, and animals with aberrant dendrite morphologies were recovered to individual plates. For details of mutations, see Table S4.

*ns302, ns303* — isolated from a screen for URX dendrite defects in genotype *ynIs78* X (*flp-8*pro:GFP)

*hmn1, hmn3, hmn4, hmn7, hmn8, hmn12* — isolated from a separate screen for URX dendrite defects in genotype *ynIs78* X (*flp-8*pro:GFP)

*hmn147, hmn159* — isolated from a screen for BAG dendrite defects in genotype *oyIs82* X (*flp-17*pro:GFP)

### Genetic mapping of mutations

With standard linkage mapping and SNP analysis, we mapped *ns303* to an interval between 18.57 Mb and 18.70 Mb on chromosome V. Genomic sequence analysis identified a mutation (W8stop) in the previously uncharacterized gene *Y51A2D.15*, which we named *grdn-1*. As no fosmid covering this region was available, we confirmed this as the causative mutation by rescuing the URX dendrite extension phenotype with a GRDN-1a cDNA driven by a ∼4.5 kb *grdn-1* promoter (see Fig. 7A; primer sequences used to amplify this promoter are in Table S3). Additional *grdn-1* alleles from the screens were identified by non-complementation to *ns303* and confirmed by Sanger sequencing.

*hmn3* was mapped to a ∼1 Mb interval on chromosome IV containing *sax-7* by a one-step whole-genome-sequencing and SNP mapping strategy (Doitsidou et al., 2010). Sequence analysis identified a candidate mutation in *sax-7,* and we confirmed this mutation by rescuing the URX dendrite extension phenotype in *hmn3* with the WRM0631cG07 fosmid and by non-complementation to the previously described *sax-7(ky146)* allele (Zallen et al., 1999). Additional *sax-7* alleles were identified by non-complementation to *hmn3* and confirmed by Sanger sequencing.

### Conventional light microscopy and image processing

Larval animals were immobilized with sodium azide, mounted on 2% agar or agarose pads, and covered with a No. 1.5 coverslip. To image embryos, adult hermaphrodites were dissected to release early-stage embryos, which were then mounted in water on slides with 2% agarose pads and covered with a No. 1.5 coverslip. Young embryos (prior to twitching, ∼1.5-fold) were imaged live, but older embryos were arrested to prevent movement. Embryo arrest was accomplished by applying 15-50 1-sec pulses of 405nm excitation light; this treatment reliably caused embryo twitching movements to cease within 20 min without morphological damage to the neurons of interest.

Image stacks were collected on a DeltaVision Core deconvolution imaging system (Applied Precision) with the InsightSSI light source; UApo 40x/1.35 NA oil immersion objective, PlanApo 60x/1.42 NA oil immersion objective, or UPlanSApo 100x/1.40 NA oil immersion objective (Olympus); the standard DeltaVision live cell excitation and emission filter set; and a Photometrics CoolSnap HQ2 CCD camera (Roper Scientific). Image stacks were acquired and deconvolved with Softworx 5.5 (Applied Precision).

Cross sections were generated using the Rotate3D Tool in Priism 4.4.0 (Chen et al., 1996). Maximum intensity projections were generated with contiguous optical sections in ImageJ (Fiji), then linearly adjusted for brightness in Adobe Photoshop CS5. Multicolored images were created by placing each channel in a separate false-colored screen layer in Photoshop. Figures were assembled in Adobe Illustrator CS5.

### Quantification of dendrite lengths

Dendrite lengths were measured using the segmented line tool in ImageJ (NIH). The dendrite was traced from the point where it joins the cell body to the point where it ends at the nose, then normalized by the distance from the cell body to the nose to account for variance in the size of the animal. All dendrite lengths were measured in L4 stage animals, except where noted. To assess the correlation between URX and BAG dendrite defects, ipsilateral URX and BAG dendrite lengths were quantified and R^2^ values were calculated for a linear regression model in Numbers (Apple).

### Dual-inverted selective plane illumination microscopy (diSPIM)

Light sheet microscopy of wild-type and *grdn-1(ns303)* embryos expressing the BAG and URX reporter *egl-13(3kb)*pro:GFP was performed on a diSPIM microscope (Applied Scientific Instrumentation, Eugene OR) (Kumar et al., 2014), though only a single imaging view was used for all experiments presented here. 488 and 561 nm laser lines from an Agilent laser launch were fiber-coupled into the MEMS-mirror scanhead, used to create a virtually-swept light sheet. A pair of perpendicular water-dipping, long-working distance objectives (NIR APO 40×, 0.8 NA, Cat. No. MRD07420; Nikon, Melville, NY) were used to illuminate the sample and to collect the resulting fluorescence. All laser lines were reflected with a quad-pass ZT405/488/561/640rpcv2 dichroic and emission was selected with a ZET405/488/561/635M filter (Chroma) before detection on an sCMOS camera (ORCA Flash v2.0; Hamamatsu). Optical stacks were acquired by sweeping the sheet in conjunction with the detection plane (controlled via piezo motor) through the sample. Before embryonic twitching began, when *egl-13*pro:GFP expression is dimmest, volumes of ∼65 z-slices (step size 0.75 µm) were acquired at 5 min intervals with a 150 msec exposure. Once twitching began, volumes were acquired at 5 sec intervals with a 20-40 msec exposure. For data acquisition and instrument control, we used the ASI diSPIM plugin within MicroManager (Edelstein et al., 2014).

Maximum intensity projections were generated and intensities were linearly adjusted using ImageJ (NIH). 8-bit images were saved as TIF files in an image sequence. Annotation was added using Adobe Photoshop CS5. The image sequence was then loaded into ImageJ and saved as a movie with a frame rate of 6 frames/sec.

### 3D-Structured Illumination Microscopy (3D-SIM)

To visualize the full membranous elaborations of BAG and URX, as well as ILso glia, we expressed combinations of cytoplasmic, membrane-anchored, and ciliary reporters under cell-specific markers (BAG: *flp-17*pro:GCY-9-mApple, *flp-17*pro:mApple, *flp-17*pro:myr-mApple; URX: *flp-8*pro:mApple, *flp-8*pro:myr-mApple; ILso glia: *grl-18*pro:GFP, *grl-18*pro:myr-GFP) (Hao et al., 2006; Gramstrup Petersen et al., 2013; Smith et al., 2013). We found that visualizing the ciliary “bag” of BAG required expression of the membrane-associated ciliary marker GCY-9-mApple.

L4 larvae grown at 20°C were anesthetized in 110 mM sodium azide and 20 mM levamisole in M9, then mounted with No. 1.5 coverslips on 3% agarose pads with 110 mM sodium azide. Imaging was performed on an OMX V4 Blaze microscope (GE Healthcare) equipped with three watercooled PCO.edge sCMOS cameras, 488 and 568 nm lasers, and 528/48 and 609/37 emission filters (Omega Optical). Images were acquired with a 60X/1.42 NA Plan Apochromat objective (Olympus) and a final pixel size of 80 nm. Optical stacks of ∼3-6 µm were acquired with a z-step of 125 nm and with 15 raw images per plane (three angles with five phases each). Spherical aberration was minimized using immersion oil matching (Hiraoka et al., 1990); generally, oil with a refractive index of 1.522 worked well. Superresolution images were computationally reconstructed from the raw data sets with a channel-specific, measured optical transfer function, and a Wiener filter constant of 0.001 using custom written 3D-SIM reconstruction code (T. Lambert, Harvard Medical School) based on (Gustafsson et al., 2008). Images are displayed as maximum intensity projections generated in ImageJ from contiguous optical sections, except Fig. 2B as described below. Negative intensity values were eliminated and images were saved as 8-bit files. Fig. 2B was generated using the Oblique Slicer tool in Imaris (Bitplane) to rotate the image volume in 3D. It was then linearly adjusted for brightness, captured using the Snap tool, and saved as an 8-bit file. All images were then imported into Photoshop CS5 and linearly adjusted for brightness. Multicolored images were created by placing each channel in a separate false-colored screen layer in Photoshop. Figures were assembled in Adobe Illustrator CS5.

### Mosaic analysis of GRDN-1 activity

Animals of genotype *grdn-1(ns303)* V; *ynIs78* X (*flp-8*pro:GFP) carried an unstable extrachromosomal transgene array that included the fully-rescuing *grdn-1*pro:GRDN-1a construct and *flp-8*pro:CFP. To obtain animals that were mosaic for the presence of the array without bias with regard to the presence of the array in URX, we selected animals in which one URX dendrite was full length and one was short, scored using only the integrated GFP marker on a fluorescence-equipped dissecting stereomicroscope. Each mosaic animal was then mounted on a slide and imaged as described above (“Conventional Microscopy and Image Processing”), with each URX neuron scored for the presence or absence of the array as indicated by the expression of *flp-8*pro:CFP.

### Quantification of GRDN-1 puncta and URX+Glia GRASP

To quantify the localization of GRDN-1 puncta relative to the URX dendrite (Fig. 7H), we imaged L1 stage animals expressing *flp-8*pro:CFP, *itx-1*pro:YFP-GRDN-1, and *itx-1*pro:myr-mCherry. A segmented line was drawn in ImageJ along the URX dendrite, from the point closest to the nose until the point where it joins the cell body. The beginning and end points of the URX dendrite were measured. When a punctum of glial YFP-GRDN-1 was directly adjacent to the URX dendrite in the x-, y-, or z-dimension, it was scored as an adjacent punctum and its position along the URX dendrite was recorded.

To quantify the extent of URX+Glia GRASP and the relative localization of GRDN-1 (Fig. 7K), we used a similar approach. However, because we could not concurrently label the URX dendrite, we could not measure GRDN-1 puncta associated with URX posterior to the GRASP signal and we could not be certain that all GRDN-1 puncta at the nose tip were associated with URX. Therefore, we measured and plotted all puncta at the nose tip, as they were typically in the same z-plane as puncta which clearly associated with the GRASP signal; this causes a discrepancy in the number of puncta at the nose between Fig. 7I and Figure 7K.

### Statistical analysis

The data were initially recorded in Microsoft Excel or Apple Numbers using the methods described above, and statistical analysis was performed using R version 3.2.3 (https://www.r-project.org/) or RStudio version 1.0.143 (https://www.rstudio.com/).

## SUPPLEMENTARY DATA

**Fig. S1.**
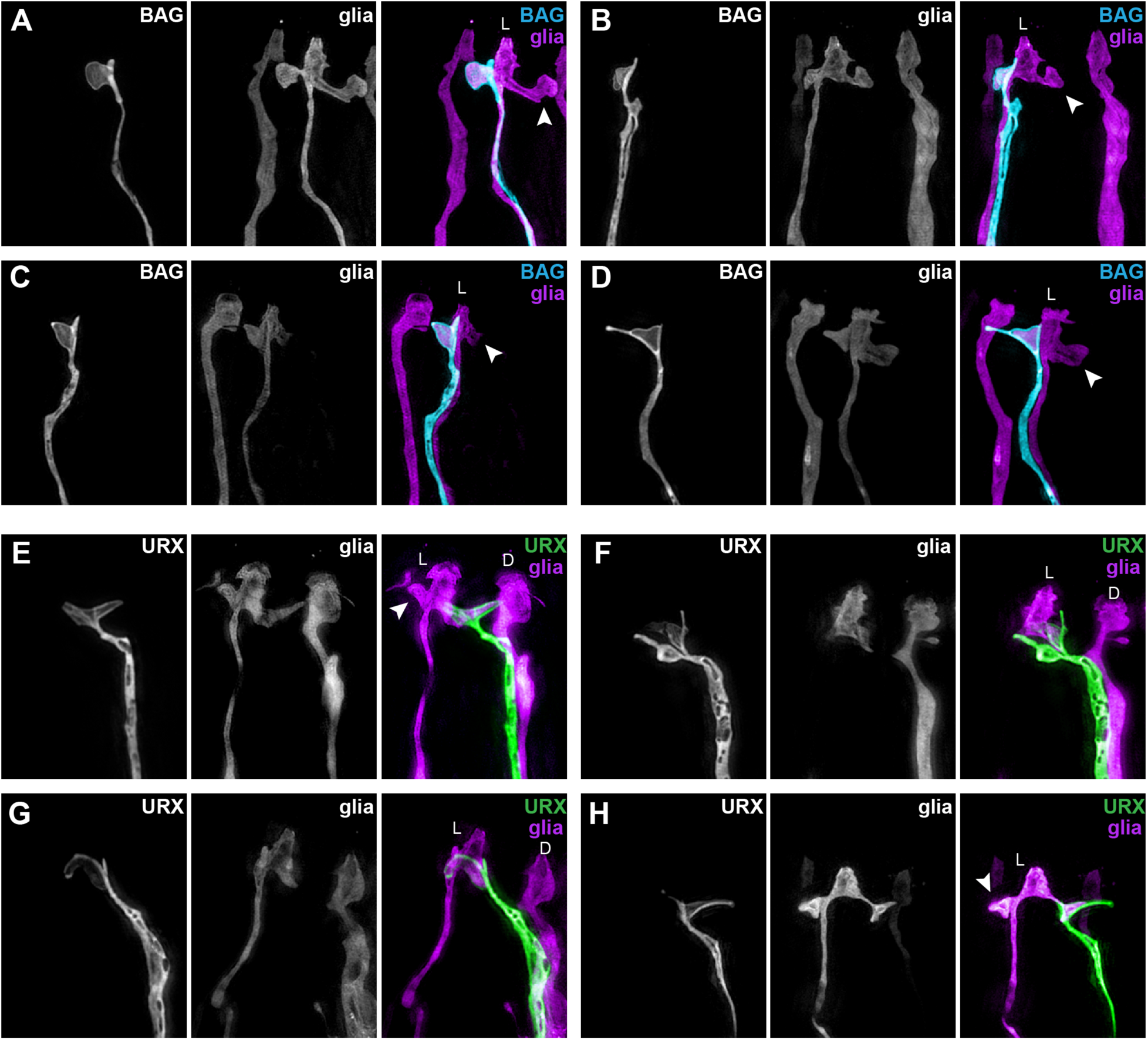
**BAG and URX dendrites form specialized contacts with a single glial cell.** Related to Fig. 1. Examples showing variability in morphology of the dendrite-glia contact across individuals. Single-wavelength and merged superresolution images of BAG (A-D, blue, *flp-17*pro) or URX (E-H, green, *flp-8*pro) and ILso glia (purple, *grl-18*pro) acquired with structured illumination microscopy. In addition to the glial protrusion that is contacted by the labeled neuron, a second glial protrusion (arrowhead) likely represents the site of contact with the unlabeled neuron (A-D, URX unlabeled; E-H, BAG unlabeled). In two examples (F, G) the second glial protrusion is hidden from view due to the orientation of the animal. The glial process in F and the dorsal glial cell in H are not visible due to these cells extending outside the optical stack. L, lateral. D, dorsal.

**Fig. S2.**
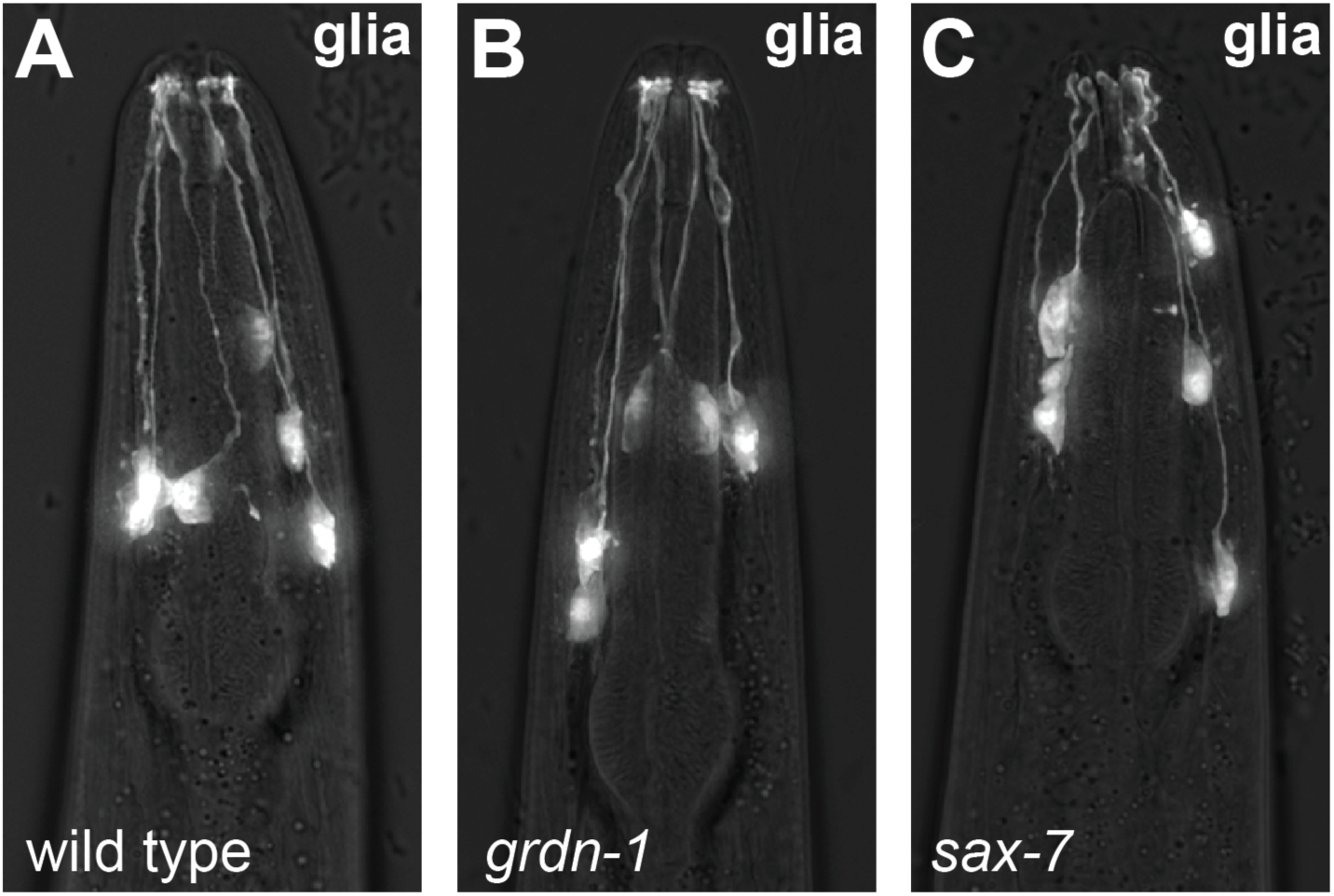
**Glial morphology in *grdn-1* and *sax-7* mutants.** Related to Fig. 3. (A) Wild-type, (B) *grdn-1*, and (C) *sax-7* animals expressing the ILso marker *grl-18*pro. Glial processes extend normally to the nose. Some cell body positioning defects are apparent in *sax-7* animals.

**Fig. S3.**
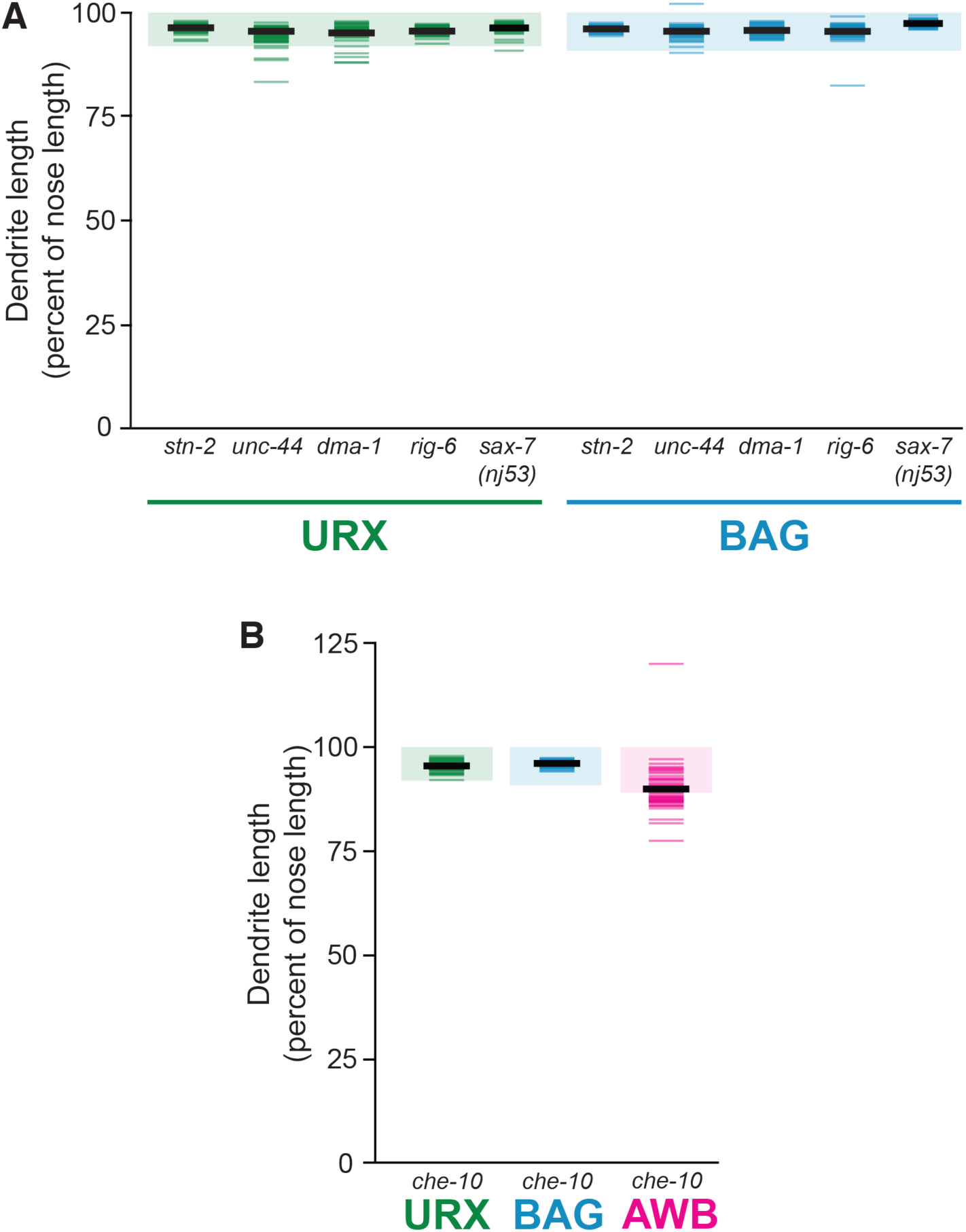
**Factors that act with SAX-7 and GRDN-1 in neuronal development are not required for URX or BAG dendrite extension.**Related to Fig. 3. Quantification of URX (*flp-8*pro, green), BAG (*flp-17*pro, blue), and AWB (amphid neuron, *str-1pro*, pink) dendrite lengths, expressed as a percentage of the distance from cell body to nose. Colored bars represent individual dendrites (n≥48 per genotype); black bars represent population averages; shaded regions represent wild-type mean ± 5 s.d. for each neuron type, which we define as “full length”. (A) Quantification of *stn-2(tm1869), unc-44(e362), dma-1(wy686), rig-6(ok1589),* and *sax-7(nj53)* dendrite lengths. SAX-7 physically interacts with UNC-44, STN-2, DMA-1, and RIG-6 to affect neuronal development, but these factors do not strongly affect URX or BAG dendrite lengths. *sax-7(nj53)* specifically affects the SAX-7L isoform, which has different adhesive properties than the SAX-7S isoform used for rescue experiments (Bénard et al., 2012). In *unc-44* mutants, URX and BAG dendrites often exhibited abnormal branching, and one BAG dendrite exhibited overgrowth. *rig-6* mutants also showed other, low-penetrance morphological defects in URX and BAG including ectopic dendrite branching and cell body positioning defects. (B) Quantification of *che-10(e1809)* dendrite lengths. GRDN-1 acts through CHE-10 in cilia development (Nechipurenko et al., 2016) and loss of CHE-10 causes dendrite extension defects in some amphid neurons, but not in URX or BAG. One AWB dendrite exhibited overgrowth.

**Fig. S4.**
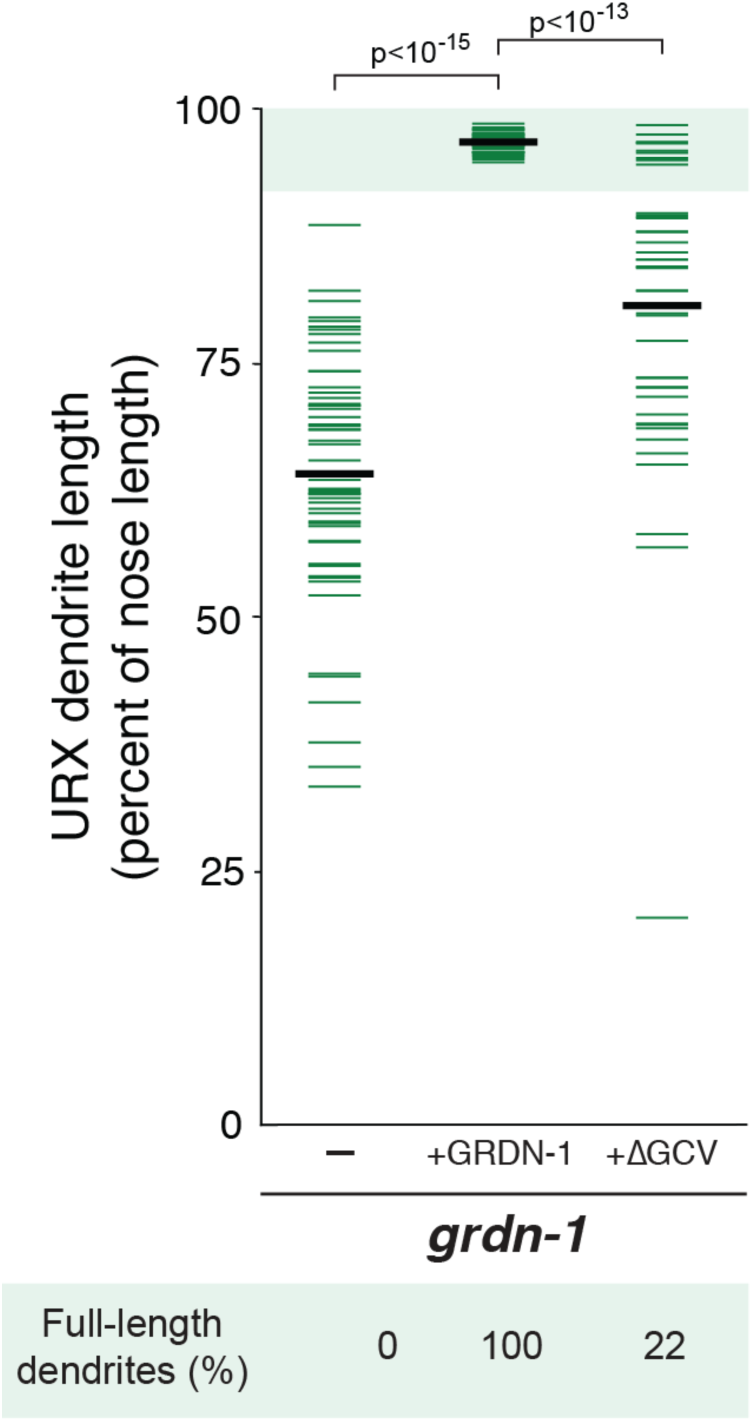
**The GRDN-1a isoform promotes dendrite extension, and requires its PDZ-binding motif for full activity.**Related to Fig. 3. Quantification of full-length URX dendrites in wild-type or *grdn-1(ns303)* animals bearing no transgene (–) or a transgene composed of a *grdn-1* promoter sequence driving expression of full-length GRDN-1a cDNA (+GRDN-1) or GRDN-1a cDNA lacking its carboxy-terminal PDZ-binding motif (+ΔGCV). Colored bars represent individual dendrites (n≥44 per genotype); black bars represent population averages. Shaded region represents wild-type mean ± 5 s.d. and the percentage of dendrites in this range (“full length dendrites”) is indicated below the plot. p-values, Wilcoxon Rank Sum test.

**Fig. S5.**
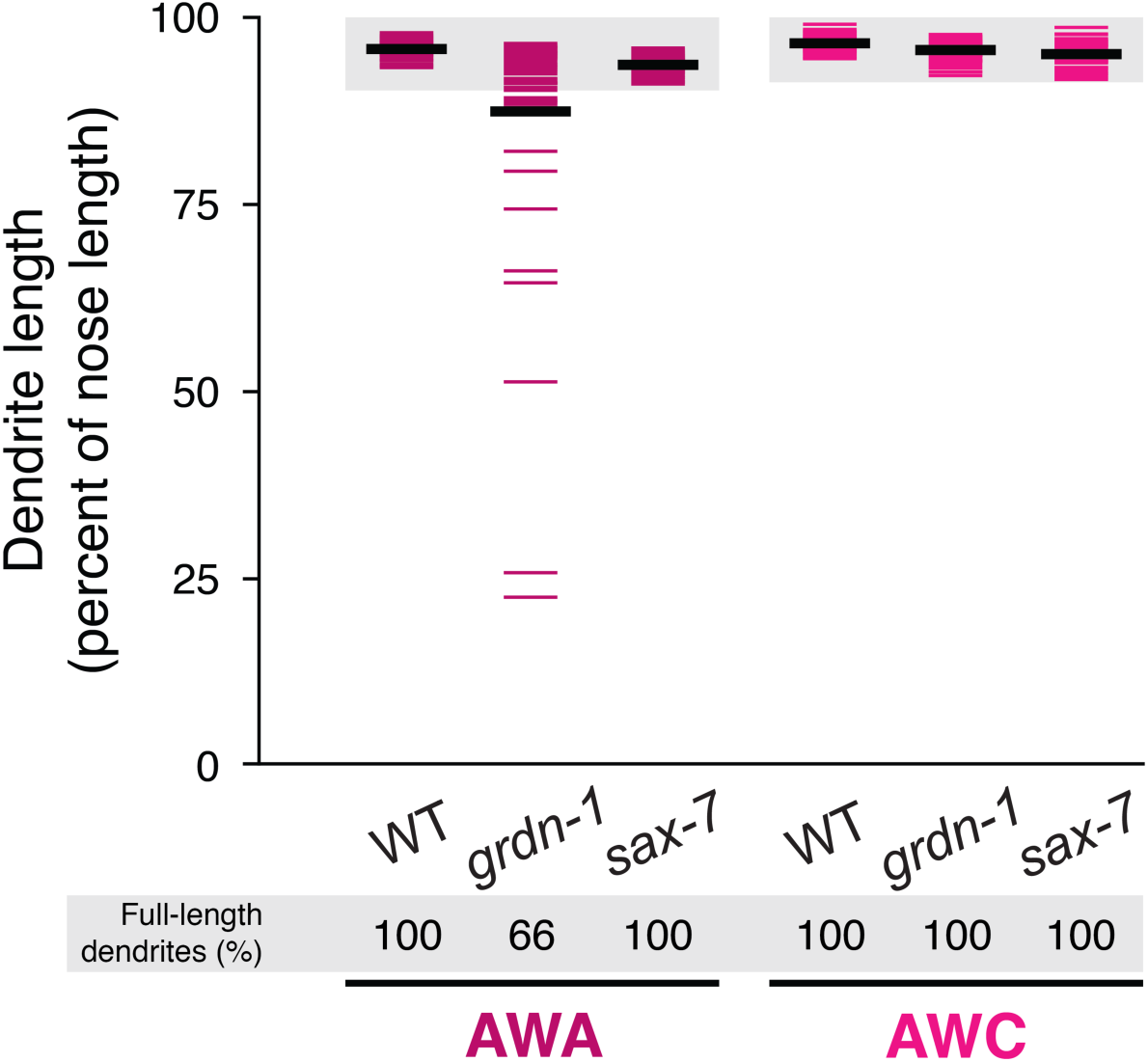
**Dendrite extension defects in amphid neurons.** Related to Fig. 4. Dendrite extension defects are observed in *grdn-1* mutants for the amphid neurons AWB (Fig. 4) and AWA, but not AWC, and these defects are not present in *sax-7* mutants. Quantification of AWA (*gpa-4Δ*pro) and AWC (*odr-1*pro) dendrite lengths in wild-type, *grdn-1* and *sax-7* animals, expressed as a percentage of the distance from cell body to nose. Colored bars represent individual dendrites (n≥50 per genotype); black bars represent population averages. Shaded region represents wild-type mean ± 5 s.d. and the percentage of dendrites in this range (“full length dendrites”) is indicated below the plot.

**Fig. S6.**
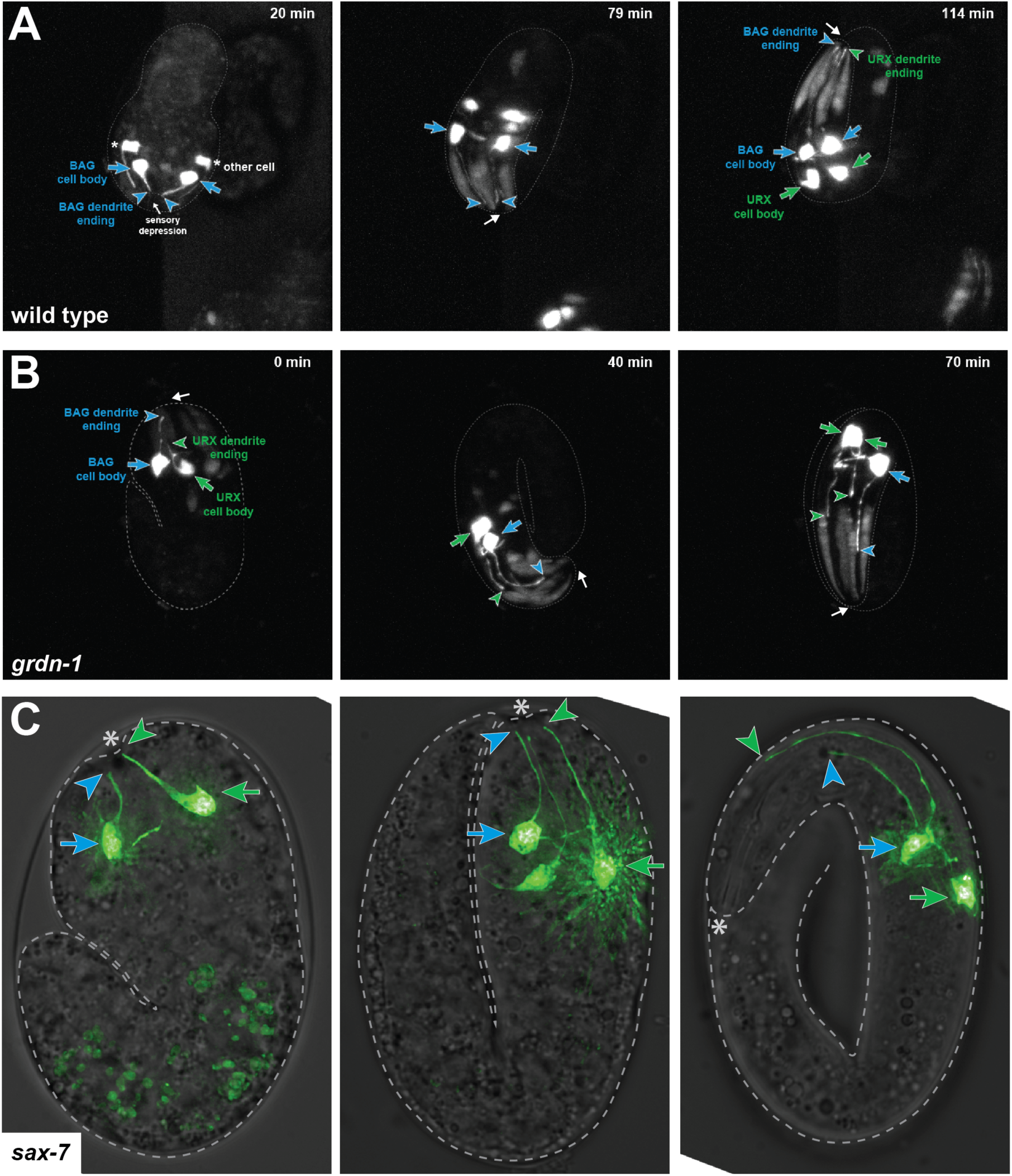
**URX and BAG dendrite length defects arise during embryo elongation.** Related to Fig. 5 and Movies S1 and S2. (A) In a wild-type embryo, BAG and URX extend short dendrites that contact the presumptive nose and appear to extend by stretch during embryo elongation. (B) In a *grdn-1* embryo, dendrites detach from the nose and fail to extend further. Time-lapse movies (Movies S1 and S2) of embryos expressing *egl-13*pro:GFP were acquired using dual-inverted selective plane illumination microscopy (diSPIM), and selected frames are shown. Time stamp is relative to start of movie, not developmental stage. At the beginning of the movies, the embryo in (A) is at a developmental stage ∼40 min earlier than in (B). White arrows indicate the nose. Colored arrows and arrowheads indicate cell bodies and dendrite endings, respectively (BAG, blue; URX, green). Approximate outline of each embryo is drawn. Changes in orientation of the animal are due to rapid twitching and writhing within the eggshell. The marker is also strongly expressed in an additional neuron (asterisk) and more weakly in elongated cells in the anterior. (C) URX and BAG dendrite extension in *sax-7* embryos at 1.5-fold, 2-fold, and pretzel stages. URX and BAG cell bodies and dendrite endings are marked with arrows and arrowheads, respectively (URX, green; BAG, blue). Asterisk, embryo nose. Dotted line, outline of embryo.

**Fig. S7.**
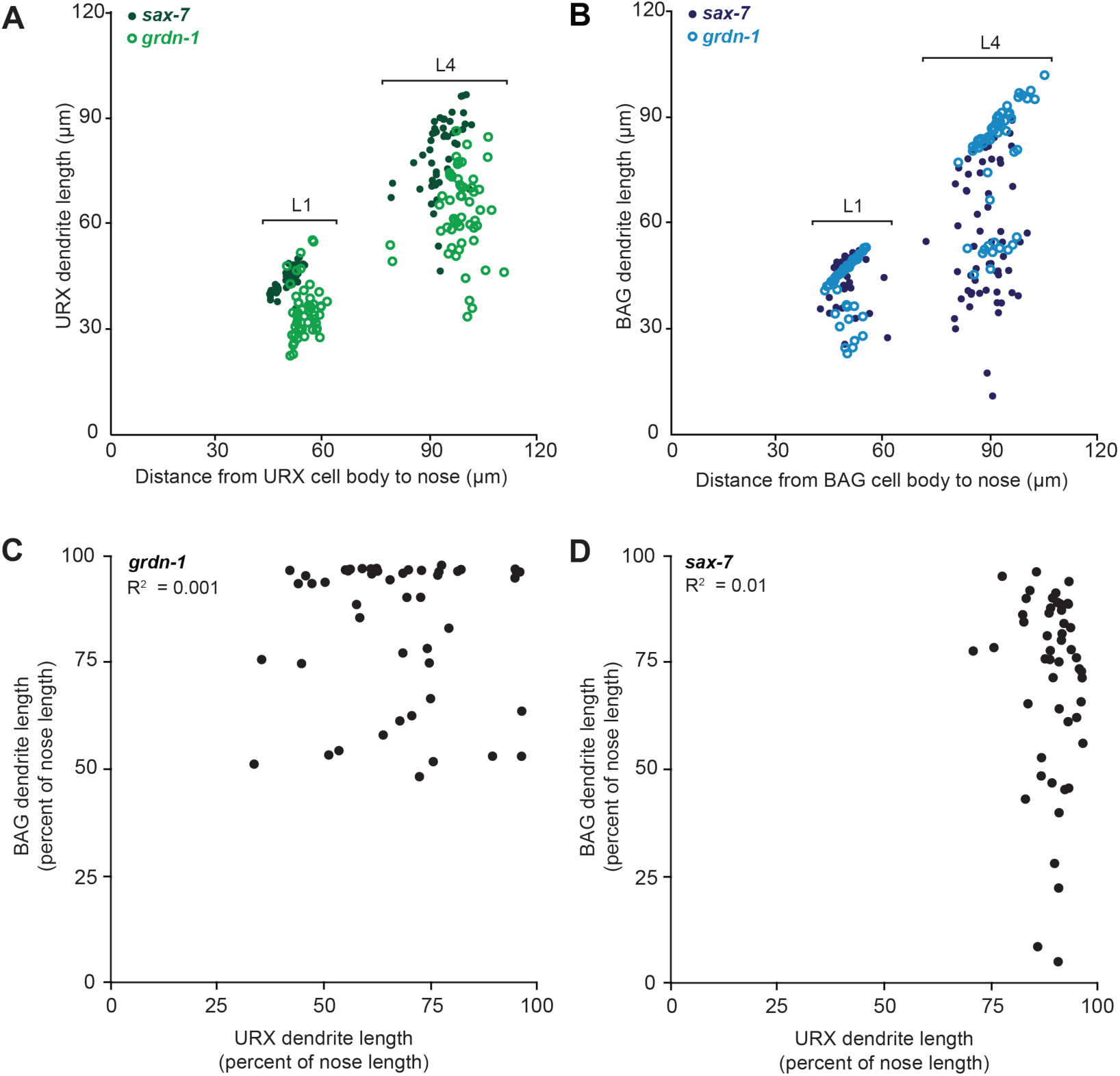
**Shortened URX and BAG dendrites keep pace with larval growth and are not correlated in length.** Related to Fig. 5. (A, B) URX and BAG dendrite lengths increase with larval growth. (A) URX and (B) BAG dendrite lengths in *sax-7* (closed circles) and *grdn-1* (open circles) animals were scored at the first (L1) and fourth (L4) larval stages and plotted as a function of the distance from the cell body to the nose. Populations of L1 and L4 animals were selected by overall morphology on a dissecting stereomicroscope. As animals grow from L1 to L4 (∼48 h), the length of the head (“Distance from cell body to nose”) approximately doubles from ∼40-60 µm to ∼80-120 µm. During this period, the lengths of the shortened URX and BAG dendrites increase similarly. n≥49 dendrites per genotype per neuron at each larval stage. (C, D) URX and BAG dendrite lengths are not correlated. Ipsilateral URX and BAG neurons were visualized simultaneously using *flp-8*pro:GFP and *flp-17*pro:mApple, respectively. Dendrite lengths were measured as a percentage of the distance from each cell body to nose. The dendrite lengths of ipsilateral URX and BAG neurons in (C) *grdn-1* and (D) *sax-7* animals are shown. URX and BAG dendrite lengths were not correlated (R^2^ = 0.001, *grdn-1*; R^2^ = 0.01, *sax-7*). n = 50 for each genotype.

**Fig. S8.**
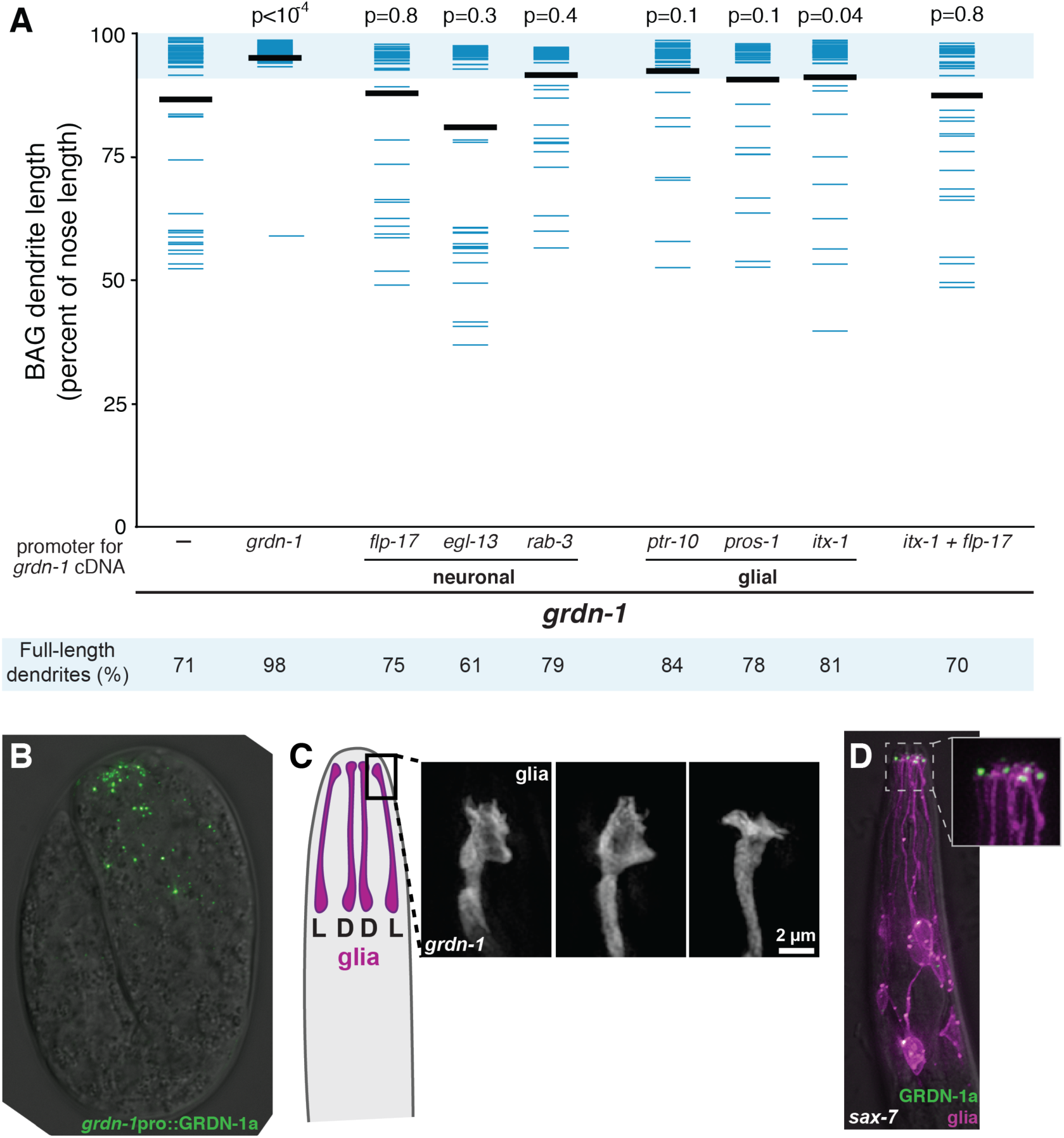
**GRDN-1 activity and localization in glia.** Related to Figure 7. (A) Transgenes containing GRDN-1a cDNA under control of the indicated promoters, or no transgene (–), were introduced into *grdn-1* animals and BAG dendrite lengths were measured as a percentage of the distance from the cell body to the nose. Expression of GRDN-1a under its endogenous promoter rescued dendrite extension defects (p<0.0001). Mild rescue using other promoters was not statistically significant (p>0.05 for all promoters shown except p=0.04 for *itx-1*pro). Colored bars represent individual dendrites (n≥40 per genotype); black bars represent population averages. Shaded region represents wild-type mean ± 5 s.d. and the percentage of dendrites in this range (“full length dendrites”) is indicated below the plot. p-values (Wilcoxon Rank Sum test) compared to *grdn-1* with no transgene (–) are at top. (B) Wild-type embryo expressing sfGFP-GRDN-1 under control of the *grdn-1* promoter. GRDN-1 is broadly expressed and localizes in puncta throughout the embryonic head. (C) Three examples of the lateral ILso glial ending in *grdn-1* animals in which both URX and BAG dendrites are short, suggesting that the elaborate protrusions characteristic of the wild-type structure are reduced or absent. Superresolution images of ILso glia (*grl-18*pro) acquired with structured illumination microscopy. Individuals with short BAG dendrites were selected for imaging; URX is always short in *grdn-1*. (D) Localization of glial-expressed GRDN-1a appeared unchanged in *sax-7* mutants. YFP-GRDN-1a (*itx-1*pro, yellow) and myristyl-mCherry (*itx-1*pro, red) were expressed together in glia. GRDN-1 localized to puncta at glial endings.

**Movie S1. URX and BAG dendrites extend by stretch during embryo elongation.** Related to Fig. 5.

<u>>>Pre-print version available at</u> http://heimanlab.com/content/2019Cebul/MovieS1.avi

Time-lapse movie of a wild-type embryo expressing *egl-13*pro:GFP, collected with dual-inverted selective plane illumination microscopy (diSPIM). BAG and URX extend short dendrites that contact the presumptive nose early in development and are then stretched to their full length during embryo elongation. The first eight frames were taken at 5 min intervals (40 min total, before the onset of embryo twitching), while the remaining frames were taken at 5 sec intervals to allow tracking of cells during rapid embryo movement. The movie is shown at a frame rate of 6 frames/sec. Selected frames have been annotated. Time stamp indicates minutes relative to start of movie. Approximate outline of embryo is drawn. Arrows and arrowheads indicate cell bodies and dendrite endings, respectively (BAG, blue; URX, green). The marker is also strongly expressed in an additional neuron (asterisk) and more weakly in elongated cells in the anterior.

**Movie S2. URX and BAG dendrite defects in *grdn-1* mutants arise during embryo elongation.** Related to Fig. 5.

<u>>>Pre-print version available at</u> http://heimanlab.com/content/2019Cebul/MovieS2.avi

Time-lapse movie of a *grdn-1* mutant embryo expressing *egl-13*pro:GFP, collected with dual-inverted selective plane illumination microscopy (diSPIM). BAG and URX extend short dendrites that contact the presumptive nose early in development, but then detach from the nose and fail to extend. Frames were taken at 5-sec intervals, and the movie is shown at a frame rate of 6 frames/sec. Selected frames have been annotated. Time stamp indicates minutes relative to start of movie. Approximate outline of embryo is drawn. Arrows and arrowheads indicate cell bodies and dendrite endings, respectively (BAG, blue; URX, green). The marker is also strongly expressed in an additional neuron (asterisk) and more weakly in elongated cells in the anterior.

**Table S1.**
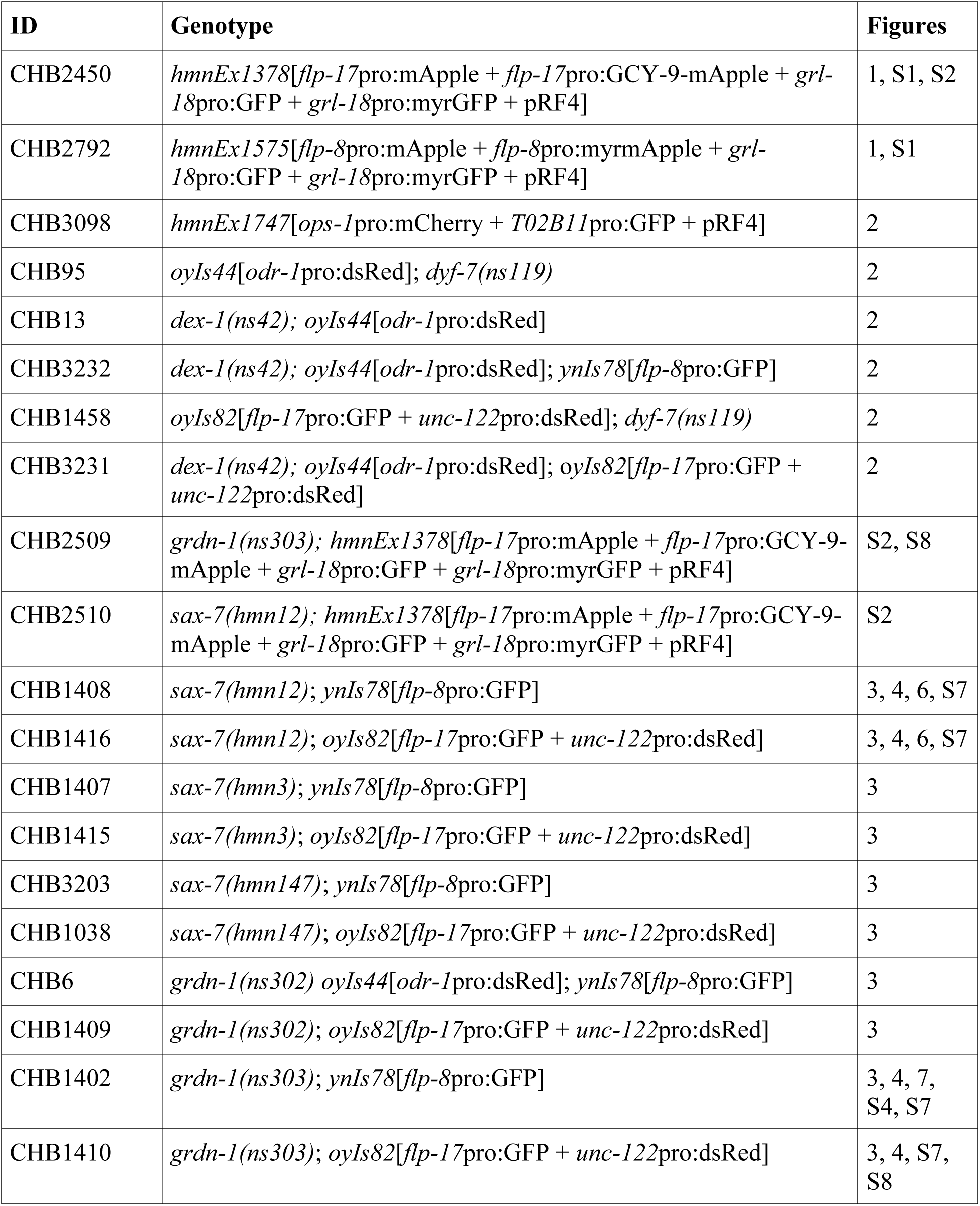
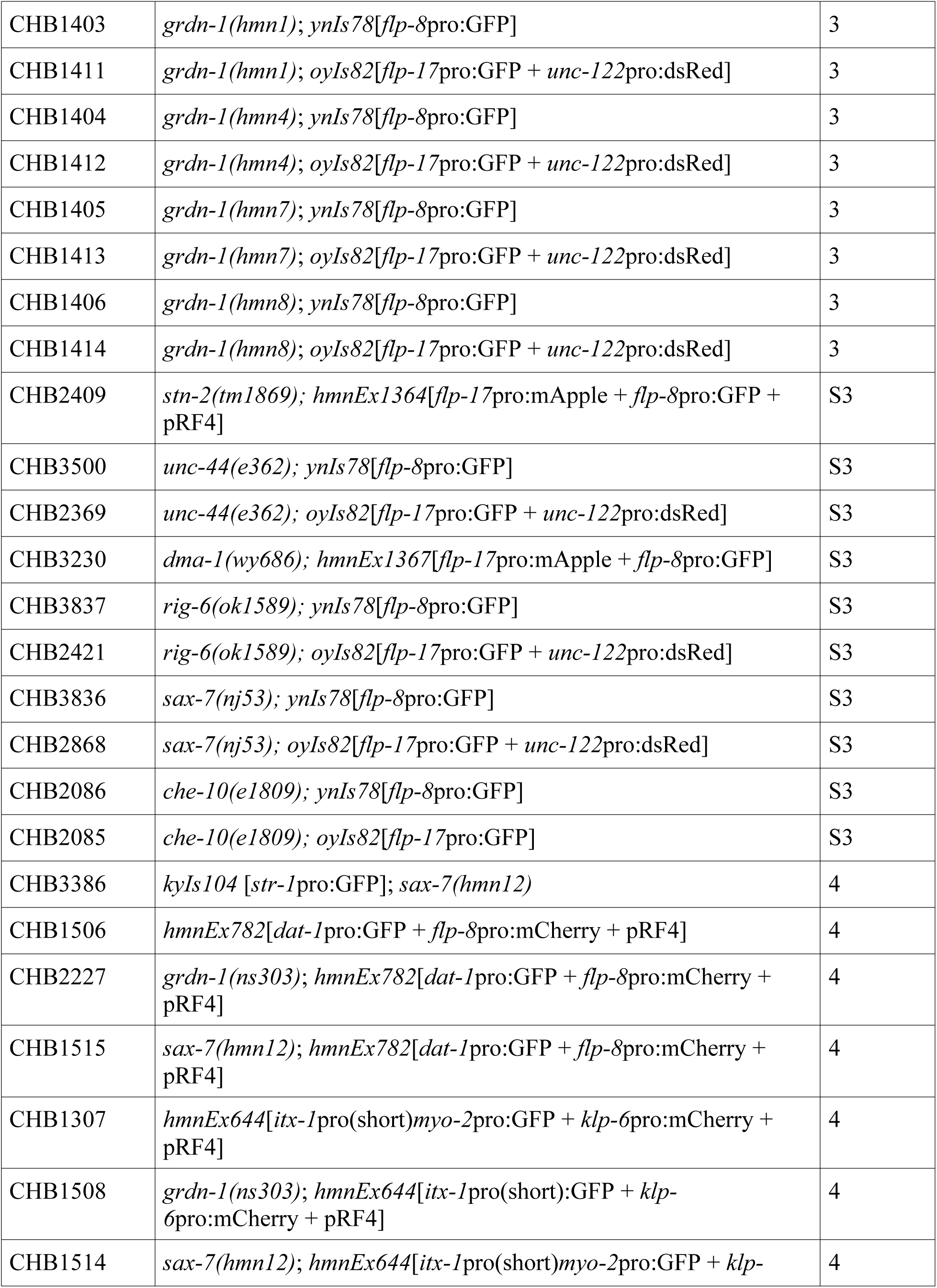
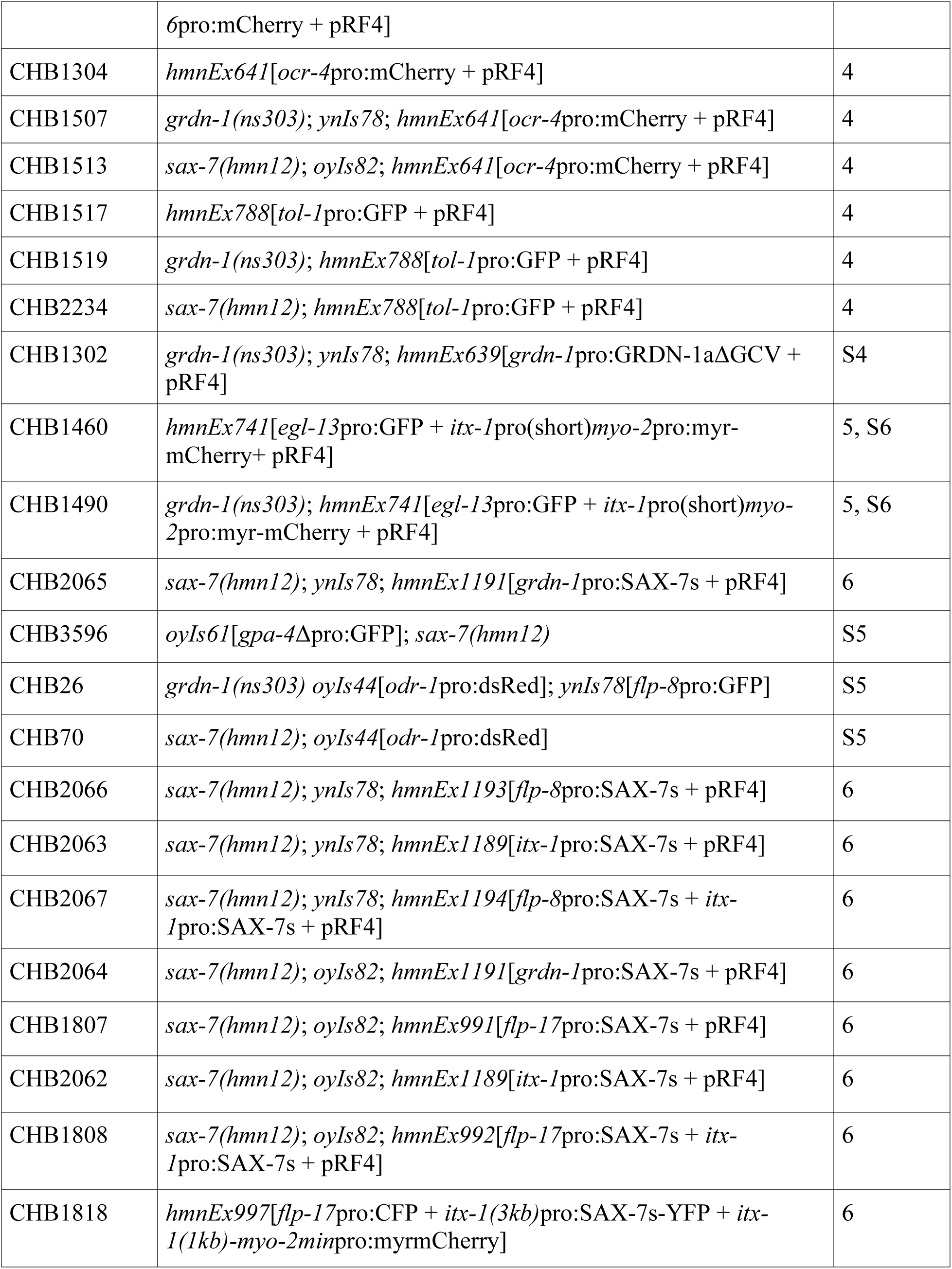
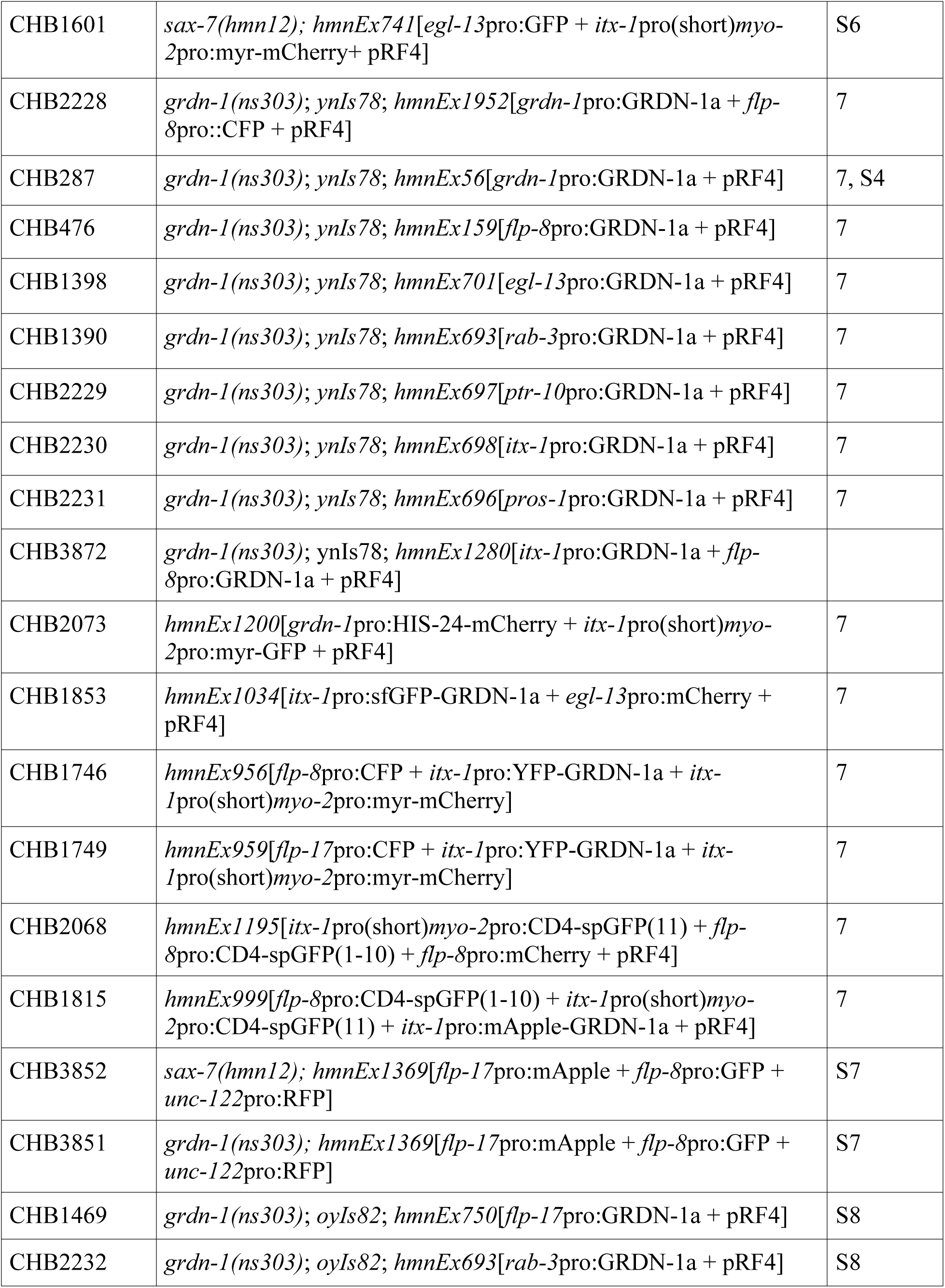
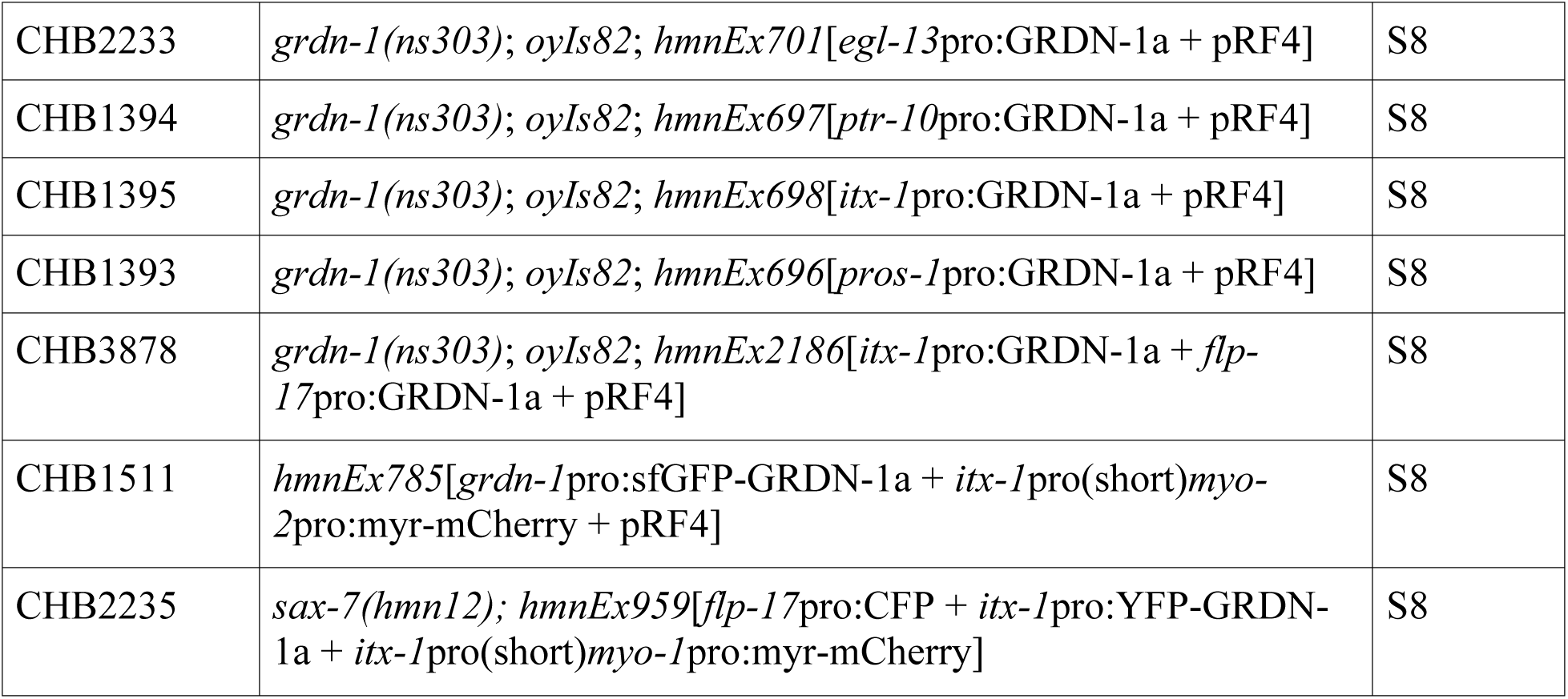
**Strains.** Related to all figures. **Strains generated for this study**

**Strains generated in previous studies.** Related to Figures 2-7 and Figures S3, S4, S5, S8.

| **ID** | **Genotype** | **Figure** | **Source/Reference** |
| --- | --- | --- | --- |
| CHB4 | *oyIs44[odr-1pro:dsRed]* | 2, S5 | Lanjuin et al., 2003<br>(as PY2417) |
| CHB1738 | *oyIs44[odr-1pro:dsRed]; dyf-7(ns119); hmnEx952[flp-8pro:CFP + tol-1pro:YFP]* | 2 | Low and Williams et al., 2019 |
| PY6342 | *che-10(e1809); kyls104[str-1pro:GFP]* | S3 | Nechipurenko et al., 2016 |
| PY1089 | *kyls104[str-1pro:GFP]* | 4 | Troemel et al., 1997 |
| PY8658 | *grdn-1(ns303); kyls104[str-1pro:GFP]* | 4 | Nechipurenko et al., 2016 |
| PY6101 | *oyIs61[gpa-4Δpro:GFP]* | S5 | Olivier-Mason et al., 2013 |
| PY8836 | *oyIs61[gpa-4Δpro:GFP]; grdn-1(ns303)* | S5 | Nechipurenko et al., 2016 |
| CHB1929 | *ynIs78[flp-8pro:GFP]* | 2, 3, 4, 6, 7, S4 | Kim and Li, 2004<br>(as NY2078) |
| CHB851 | *oyIs82[flp-17pro:GFP, unc-122pro:dsRed]* | 2, 3, 4, 6, S8 | Astrid Cornils and Piali Sengupta (as PY8503) |

**Table S2.**
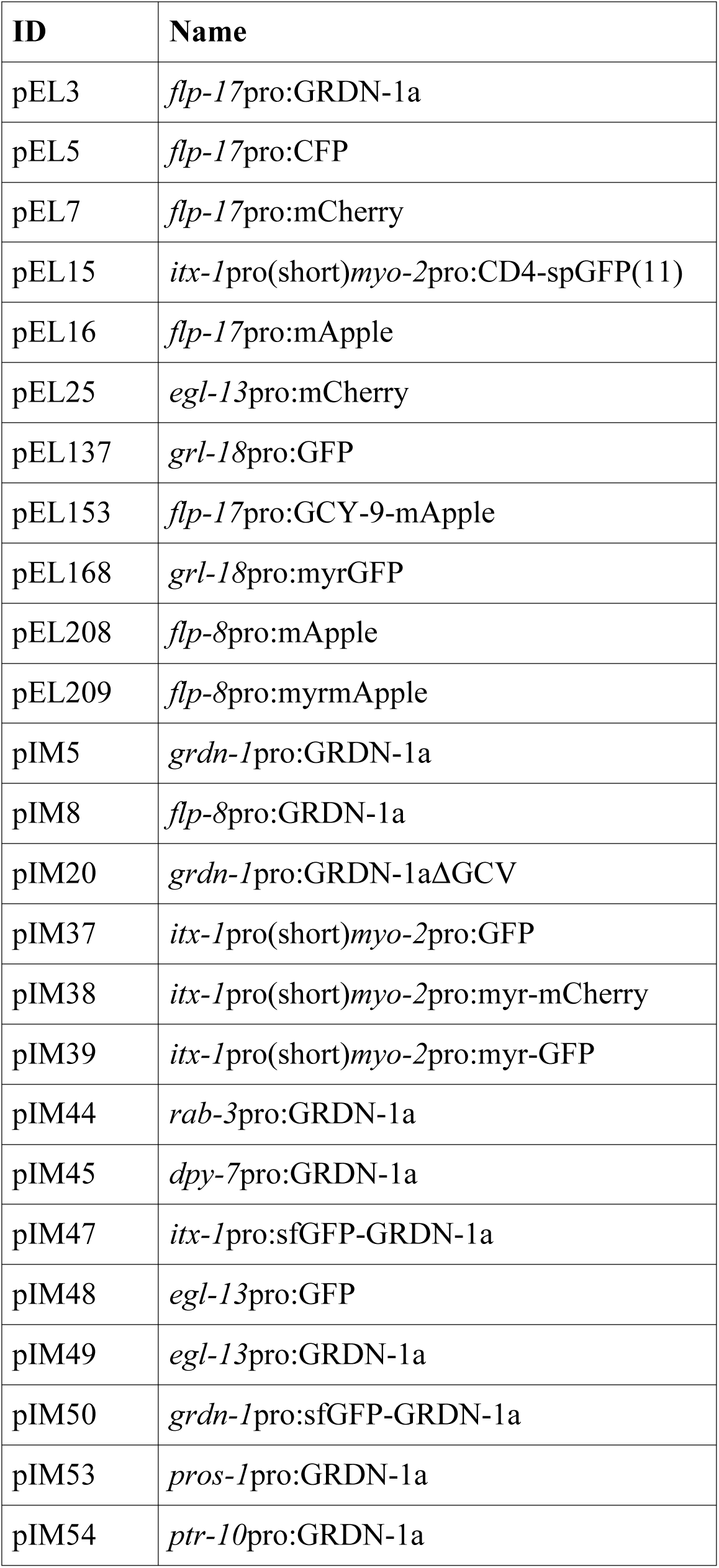
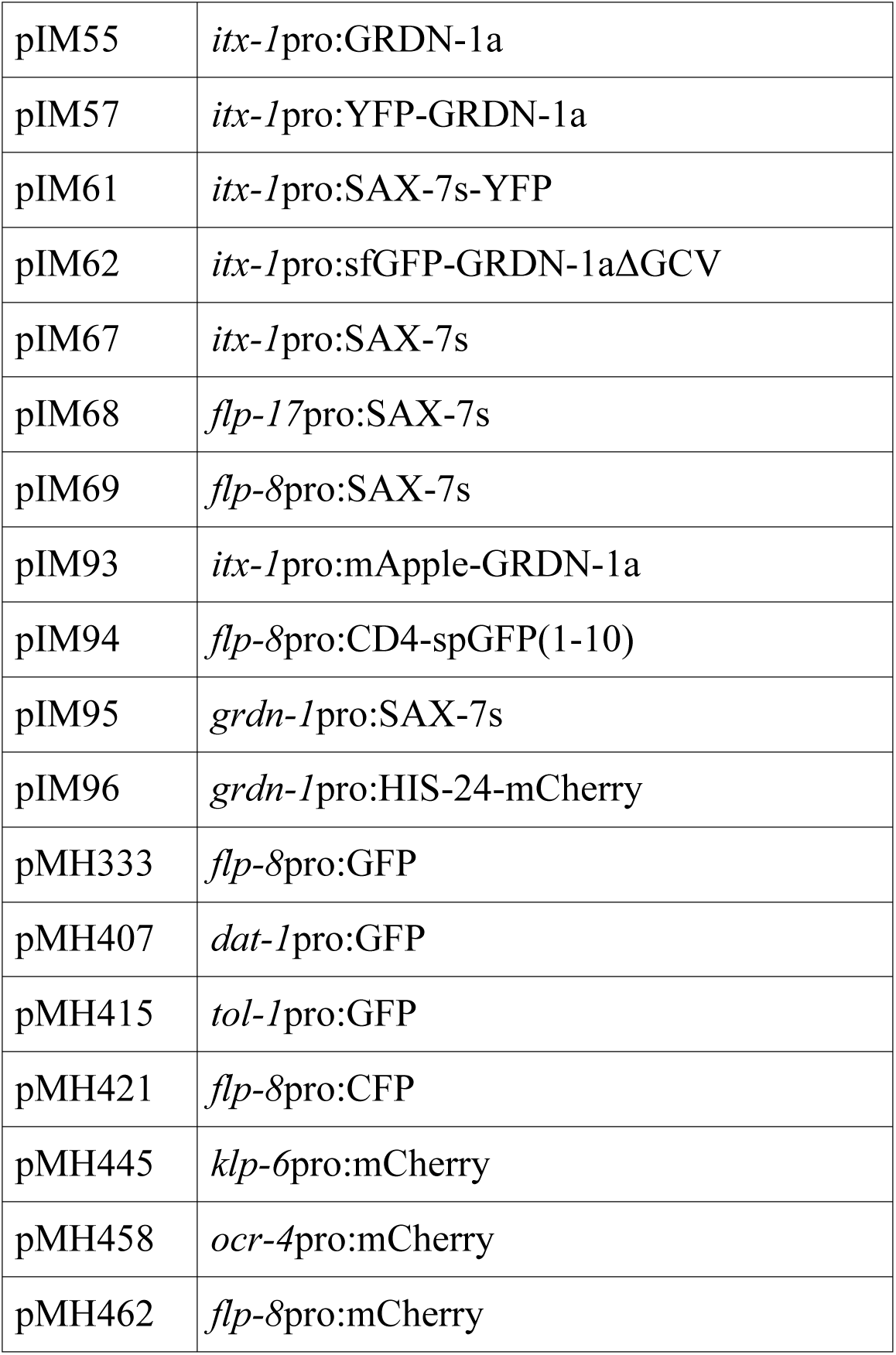
**Plasmids generated for this study.**

**Table S3.**
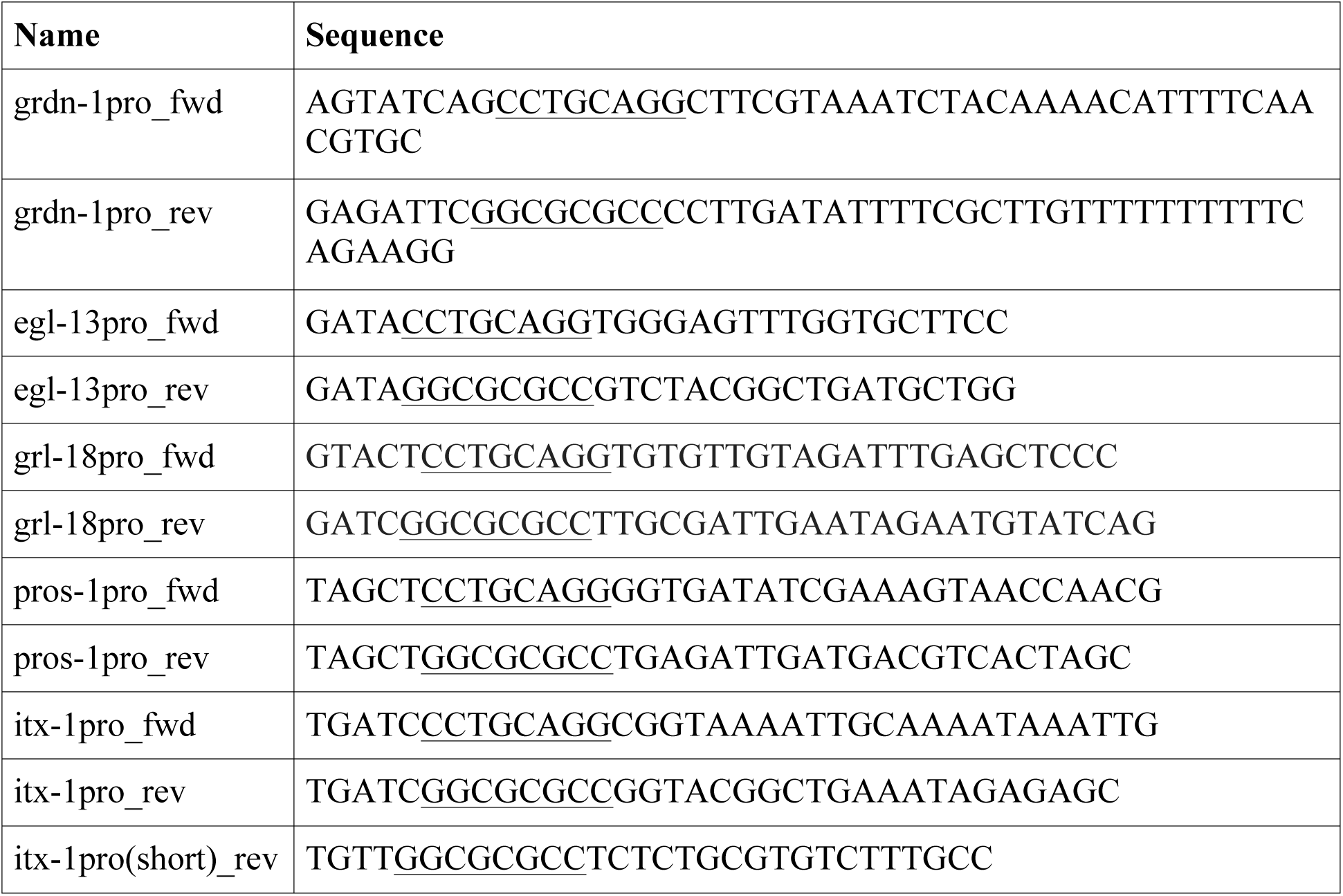
**Primers** Primers of general interest

**Table S4.**
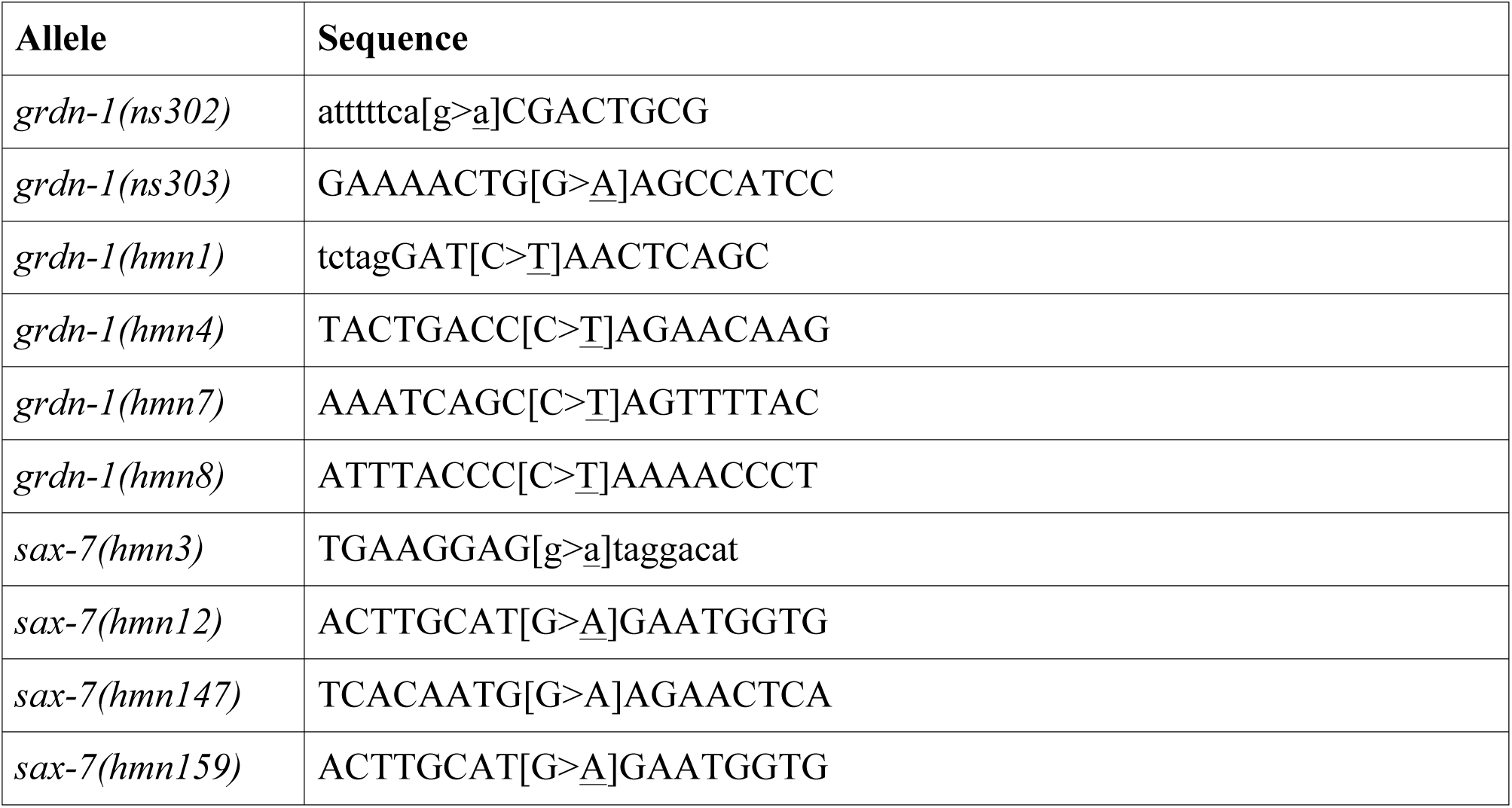
**Mutant alleles.** Related to Figure 3. Alleles isolated in this study. Substitutions are bracketed with the mutant sequence underlined. Uppercase corresponds to predicted exons; lowercase to predicted introns.

## Notes

http://heimanlab.com/content/2019Cebul/

